# The ubiquitin ligase Hecw controls oogenesis and neuronal homeostasis by promoting the liquid state of ribonucleoprotein particles

**DOI:** 10.1101/2020.05.30.124933

**Authors:** Valentina Fajner, Fabio Giavazzi, Simona Sala, Amanda Oldani, Emanuele Martini, Francesco Napoletano, Dario Parazzoli, Roberto Cerbino, Elena Maspero, Thomas Vaccari, Simona Polo

## Abstract

Specialised ribonucleoprotein (RNP) granules are a hallmark of germ cells. Among their main function is the spatial and temporal modulation of the activity of specific mRNA transcripts that allow specification of primary embryonic axes. While RNPs composition and role are well established, their regulation is poorly defined. Here, we demonstrate that Hecw, a newly identified *Drosophila* ubiquitin ligase, is a key modulator of RNPs in oogenesis. Loss of Hecw activity results in the formation of enlarged granules that transition from a liquid to a gel-like state. At the molecular level, Hecw depletion leads to reduced ubiquitination and activity of the translational repressor Fmrp, resulting in premature Orb expression/recruitment in nurse cells. In addition to defective oogenesis, flies lacking Hecw show neurodegenerative traits with premature aging and climbing defects due to neuronal loss that are linked to RNPs condensation. Our findings reveal an unprecedented function of ubiquitin in modulating RNP fluidity and activity.

## INTRODUCTION

In the ubiquitination cascade, E3 ligases act as catalysts and molecular matchmakers of the reaction. They carry out the final step of covalently binding of ubiquitin (Ub) to substrates, usually to lysine (K) residues, and define the type of Ub chains attached to them, dictating targets fate. Among the HECT (Homologous to the E6-AP Carboxyl Terminus) E3s families that are present in all eukaryotes, NEDD4 is the most characterised, with one member in yeast, three in *Drosophila melanogaster* and nine in humans. NEDD4 ligases modify multiple substrates by monoubiquitination^1^ or by addition of K63-linked Ub chains^2^ to support a variety of cellular functions, such as protein trafficking, signalling regulation, or lysosomal degradation. By virtue of such multifaceted activity, the Nedd4 family of HECT ligases is known to contribute to a wide range of physiologic and pathologic processes, including immune regulation, viral infection, tumorigenesis and neurological disorders^3,4^.

In this study, we investigate the molecular, cellular and organismal functions of a previously uncharacterised single *Drosophila* ortholog of HECW1 and 2. We show that Hecw is required to support fertility and neuronal health by ubiquitinating the fragile X mental retardation protein (Fmrp), ultimately regulating translational repression of key developmental factors within ribonucleoparticles (RNPs). By characterizing the biophysical properties of Me31B-labeled germ granules, a type of membraneless RNP organelles also called P bodies or sponge bodies^5^, we demonstrate that the Hecw modulates their state of aggregation, promoting fast molecular exchange and repression activity. Our findings reveal an unexpected function for ubiquitination in maintaining the liquid state of germ granules that may be generally relevant to the functions of RNPs.

## RESULTS

### CG42797/Hecw is the *Drosophila* ortholog of HECW1 and HECW2

By searching the *Drosophila* genome, we identified a single ortholog of human HECW1 and HECW2, encoded by the uncharacterised gene CG42797, whose protein product shares co-linearity and 40% overall similarity with the corresponding human proteins. Extensive amino acid sequence identity is present in two WW domains required for substrate interaction^6^ and in the catalytic HECT domain (**Fig. 1a**; **Supplementary Fig. 1a**). Similar to the *C. elegans* Ce01588 (ref. ^7^), the *Drosophila* protein lacks the C2 lipid binding domain, which is characteristic of the NEDD4 family, suggesting that CG42797 may be unable to bind membranes^7^.

**Figure 1.**
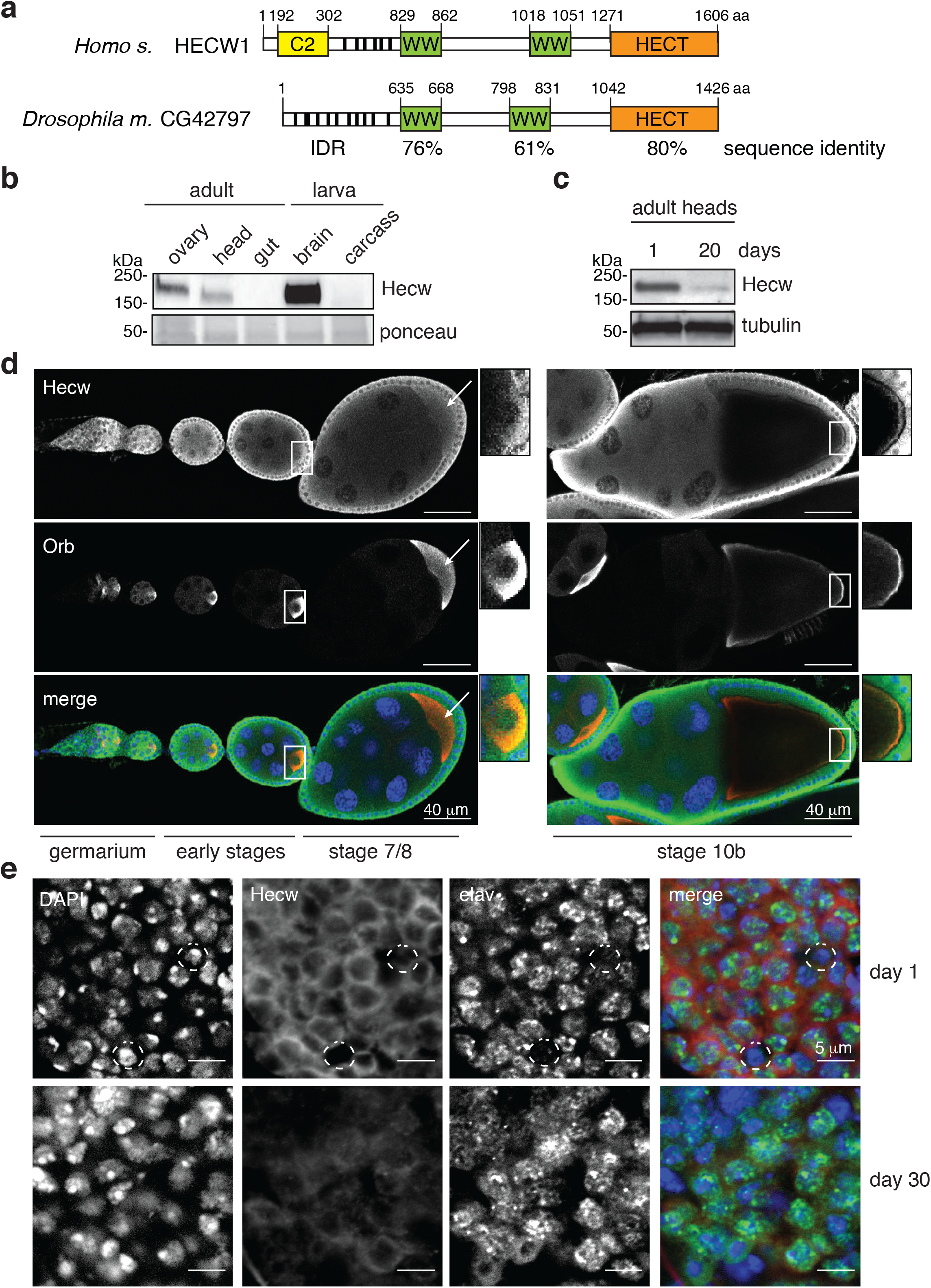
Hecw is preferentially expressed in fly gonads and central nervous system. (**a**) Schematic representation of human HECW1 and *Drosophila* CG42797 proteins. Domains shown are: C2 (Ca^2+^ dependent lipid binding) domain in yellow, WW (substrate interacting) domains in green, and HECT (catalytic) domain in orange. The fly ortholog shows no C2 domain and presents an extended intrinsically disordered region (IDR), identified by IUPred (https://iupred2a.elte.hu/). Percentage of identity is reported below the domains. (**b**) Immunoblot (IB) analysis of the indicated *Drosophila* tissues performed with the anti-Hecw antibody. Ponceau shows equal loading. (**c**) IB of adult heads of young and mature flies with the indicated antibodies. (**d**) Immunofluorescence (IF) analysis of ovarioles of 3-day-old flies performed with the indicated antibodies. Left panels: germarium and previtellogenic stages 1-7. Right panels: stage 10 egg chamber. Hecw colocalising with Orb (oocyte marker) at the posterior margin of the fly oocyte is indicated by white arrows and highlighted in the magnifications. Scale bar: 40 μm. (**e**) IF analysis of adult brains of 1-day-(upper panels) or 30-day-old flies (bottom panels) with the indicated antibodies. Neurons are marked with the anti-elav antibody and non-neuronal cells are circled in white. Scale bar: 5 μm.

To confirm that CG42797 is a catalytically active HECT ligase, we performed an *in vitro* self-ubiquitination assay. We found that CG42797 is able to ubiquitinate itself and to generate free polyubiquitin chains, whereas a mutation in the evolutionary conserved catalytic cysteine abrogates both activities (**Supplementary Fig. 1a,b**). To identify the type of Ub linkage generated by CG42797, we set up the *in vitro* reaction with K/R mutated Ub molecules and immunofluorescence experiment with specific antibodies. Results indicate that CG42797, like HECW1 (ref. ^8^) and other NEDD4 family members^9^, preferentially generates K63-specific Ub chains both *in vivo* and *in vitro* (**Supplementary Fig. 1c,d**). Based on the functional and structural similarities with its human orthologs, we named the *Drosophila* CG42797 gene ‘Hecw’.

Previous studies indicate that human HECW1 and HECW2 are preferentially expressed in neuronal tissue^10,11^. Analysis of various *Drosophila* organs by qPCR and immunoblotting, revealed a preferential expression of *Hecw* in the fly gonads and central nervous system (**Fig. 1b,c; Supplementary Fig. 1e,f**). In fly ovaries, Hecw displays a broad distribution in the cytoplasm of both somatic and germline tissues (**Fig. 1d**). Similarly, the adult brain showed cytoplasmic staining exclusively in elav-positive neuronal cells (**Fig. 1e**). Interestingly, like the components of the ubiquitin proteasome system and autophagy pathway controlling protein homeostasis^12^, also the level of Hecw expression in fly heads decreases with aging (**Fig. 1c,e**; **Supplementary Fig. 1g**).

### Loss of Hecw causes age-dependent neuronal degeneration

To study the consequences of loss of *Hecw*, we generated catalytic inactive (CI) mutants and knock out (KO) flies using the CRISPR/Cas9 system. As all the mutants and KO lines tested exhibit identical defects (**Supplementary Fig. 2**), we hereby describe *Hecw^CI^* and *Hecw^KO^* as representative examples.

Both *Hecw^CI^* and *Hecw^KO^* homozygous flies (mutant from now on) are viable and do not show macroscopic morphological defects, indicating that *Hecw* is a non-essential gene. Prompted by the reduced expression of Hecw in adult flies, we investigated the behaviour of the mutant flies during aging. To minimise the influence of genetic background, environment, nutrition and mating conditions^13,14^, we performed a lifespan assay with mixed-sex groups in standard cornmeal food. *Hecw^CI^* and *Hecw^KO^* mutant animals displayed reduced longevity with respect to isogenic control lines with a 22% and 24% decrease in median survival, respectively (**Fig. 2a,b**). Mutant lifespan reduction was even higher at 29°C (28% median survival reduction, **Supplementary Fig. 3a**).

**Figure 2.**
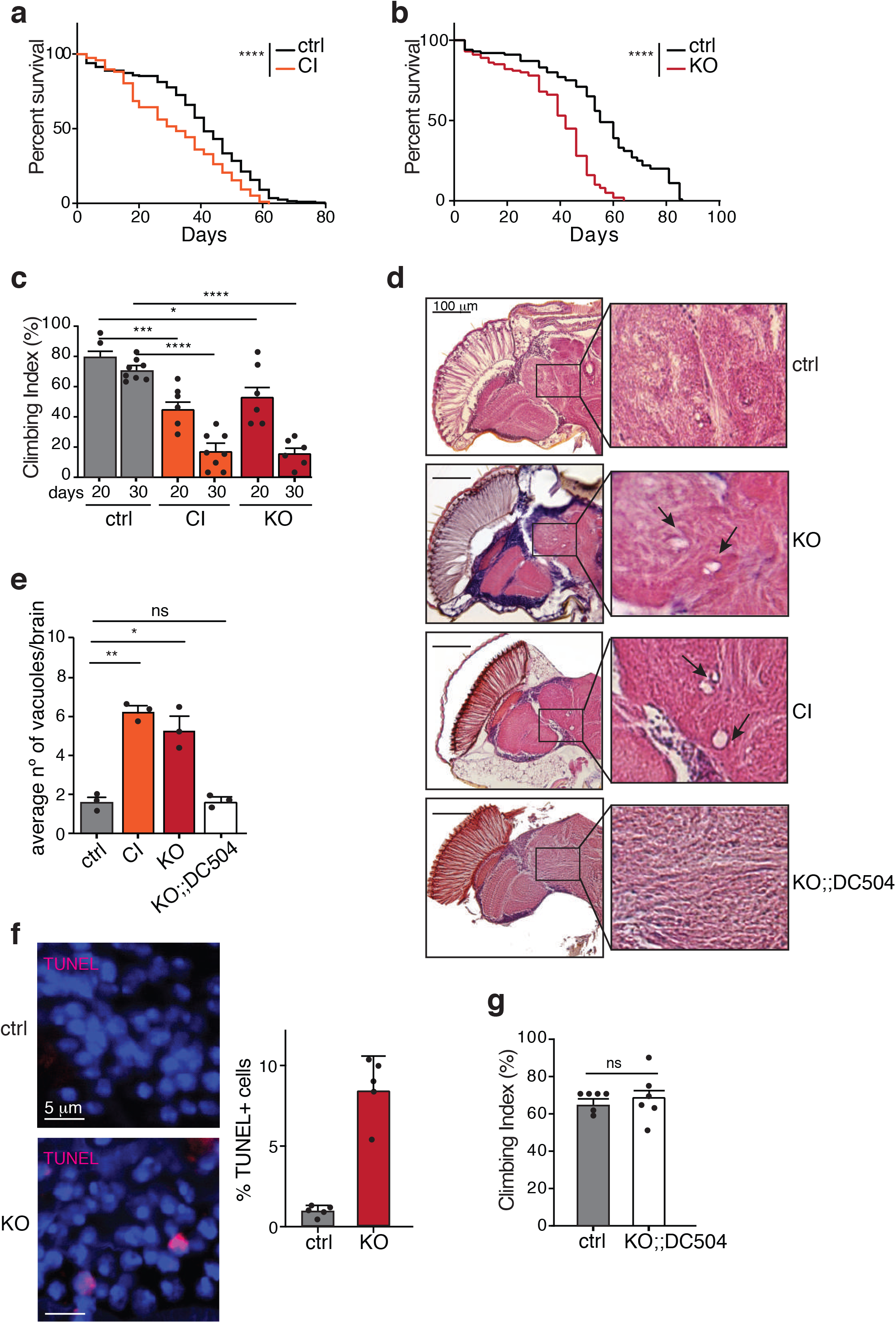
Hecw mutants display neurodegenerative phenotype. (**a,b**) Survival curve of the indicated genotypes. Percentage of survivals was calculated over 200 animals/genotype. *Hecw^CI^* (CI) (**a**) and *Hecw^KO^* (KO) flies (**b**) show a significant decrease in their lifespan compared with their control lines. **** P<0.0001 by log-rank (Mantel–Cox) test. (**c**) Motor function ability measured at the indicated time points by negative geotaxis assay. Results are expressed as mean of 8 groups of animals (*n*=8/group) ± s.e.m. * P<0.05, *** P<0.001, **** P<0.0001 by Mann Whitney test. (**d**) Frontal sections of 30-day-old fly brains of the indicated genotypes stained with H&E and examined by bright-field microscopy. Scale bar, 100 μm. Right, magnification of the central brain regions is highlighted. (**e**) Quantification of the vacuoles with diameter >2 μm in 30-day-old fly brains of the indicated genotypes, *n*=5 animals/genotype. Results are expressed as mean of three biological replicates ± s.e.m. (**f**) TUNEL staining of 30-day-old fly brains of the indicated genotypes. Right, quantification expressed as percentage of positive cells. 200-500 cells were counted for each fly, *n*=5 animals/genotype. **** P<0.0001 by two-tailed t-test. (**g**) Negative geotaxis assay performed as in c, on 20-day-old flies of the indicated genotypes. Results are expressed as mean of 8 groups of animals (*n*=8/group) ± s.e.m. NS, not significant by Mann Whitney test.

*Drosophila* displays an aging-related decline in climbing^15^ whose anticipation is considered a hallmark of neuronal dysfunction. Remarkably, in climbing assay^16^, both Hecw mutant flies displayed motor function impairment, which was already visible in 20-day-old flies and became extremely severe as the flies aged (**Fig. 2c**). Again, the phenotype was exacerbated at 29°C (**Supplementary Fig 3b**). Next, we analysed frontal brain paraffin sections and observed that mutants presented extended tissue vacuolisation (**Fig. 2d,e**), which resulted from neuronal death in the CNS (**Fig. 2f**). The size and number of vacuoles are significantly higher in mutants compared to isogenic control flies (**Fig. 2e**) and tissue vacuolisation progressively increases with fly age (**Supplementary Fig 3c**).

Importantly, defective neuronal function and morphology are rescued by the reintroduction of a wild-type copy of *Hecw* in homozygous *Hecw^KO^* animals (**Fig 2d, g**). Moreover, overexpression of Hecw with the pan-neuronal driver *elav-GAL4* causes reduction in longevity, similarly to that observed in *Hecw* mutants (**Supplementary Fig. 3d-f**), indicating that Hecw activity needs to be tightly regulated to protect neurons from premature neurodegeneration.

### Loss of Hecw leads to defective oogenesis

The high expression level of Hecw in fly gonads prompted us to assess gamete formation in mutant flies (**Fig. 1b**, **Supplementary Fig 1f**). In females, egg-laying varies with age. After a peak at day 4 post eclosion, there is a physiological decline in egg production, which is generally reduced by 50% at 40 days^17^. When compared to matched control flies, 20-day-old *Hecw* mutant females lay a significantly reduced number of eggs, (**Fig. 3a**). To determine whether the reduced egg-laying exhibited by *Hecw* mutants is accompanied by structural defects, we immunostained ovaries of 3- and 30 day-old well-fed mated wild-type and mutant flies to detect the CPEB protein Orb (oo18 RNA binding), which accumulates in the cytoplasm of the developing oocyte^18^. Already at day 3, 20% of egg chambers derived from mutant flies showed an altered number of germ cells (**Fig. 3b**). Defective egg chambers present either reduced or increased number of nurse cells and ring canals, and compound egg chambers (**Fig. 3b**, **Supplementary Fig 4a**). In addition, we detected the presence of late stage apoptotic egg chambers that are usually absent in nutrient-rich conditions (**Fig. 3b**), indicating that defective egg chambers are likely culled. Consistently, hatching rate of mutants is not significantly affected (**Supplementary Fig. 4b**). Penetrance of these defects increases with age, reaching about 40% in 30-day-old flies. Classification and quantification of the aberrant phenotypes observed are summarised in **Supplementary Table 1**.

**Figure 3.**
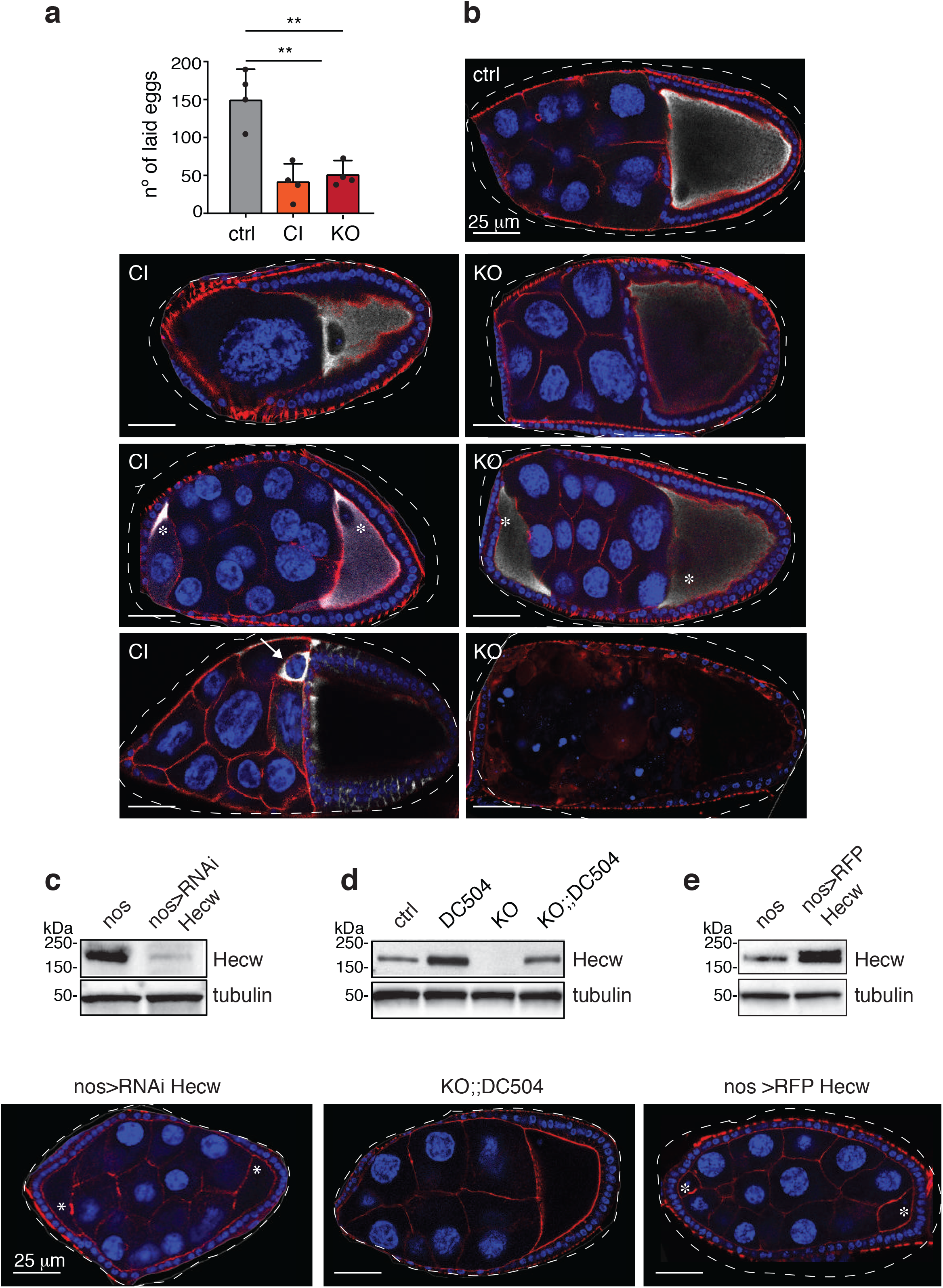
Aberrant expression of Hecw impairs proper oogenesis. (**a**) Fertility assay of the indicated genotypes. *n*=6 20-day-old females/genotype. Results of four biological replicates are expressed as mean ± SD, **P<0.01 by Mann Whitney test. (**b**) IF analysis of wild-type (ctrl, upper right panel), *Hecw^CI^* (CI, left panels) and *Hecw^KO^* (KO, right panels) egg chambers. Mutants show aberrant number of nurse cells (top panels), compound egg chambers (middle panels), oocyte misspecification and apoptotic egg chambers (bottom panels). Blue, DAPI; red, phalloidin; white, Orb; asterisks indicate oocytes in compound egg chambers; white arrow indicates an Orb-positive nurse cell. Scale bar: 25 μm. (**c, d, e**) IB and IF analysis of the indicated genotypes. Hecw depletion in the germline causes oogenesis defects, as in b, upper panel. Example of compound egg chamber in a 3-day-old Hecw-depleted fly (*nos>RNAi Hecw*) (c). The oogenesis defects are rescued in the *Hecw^KO^:DC504* genetic background (d). Ectopic expression of Hecw in the germline (*nos>RFP-Hecw*) induces oogenesis defects (e). Blue, DAPI; red, phalloidin. Asterisks indicate oocytes of the compound egg chamber. Scale bar: 25 μm. See also Supplementary Table 1 for the quantification.

To attribute these functional alterations either to the somatic or germline cells, we performed tissue-specific knock down in follicle cells (FC), using the *traffic-jam* driver (tj-Gal4), and in germline cells (GC), using the *nanos-Gal4* driver. Depletion of Hecw in GC (**Fig. 3c**) but not in FC (**Supplementary Fig. 4c**) recapitulates *Hecw^CI^* or *Hecw^KO^* mutant phenotypes, demonstrating that aberrant oogenesis defects are mostly caused by the absence of Hecw function in the germline.

The oogenesis defects of chronically Hecw-depleted flies are fully rescued by the reintroduction of a wild-type copy of *Hecw* expressed at physiological levels in the whole organism (**Supplementary Table 1**, **Fig. 3d**). As shown in neurons, unscheduled expression of Hecw is detrimental also in gonads, in which ectopic expression of Hecw leads to oogenesis aberrations similar to those observed in *Hecw^CI^* animals (**Fig. 3e, Supplementary Fig.4d**). These data indicate that germline Hecw expression is tightly regulated to support oocyte development.

### Loss of Hecw changes the aggregation state of RNPs

In developing egg chambers, Orb protein expression is usually confined to the oocyte thanks to the repressive action of Cup and Fmrp on selected mRNAs, as part of RNPs assembled in nurse cells^19–21^. Interestingly, we observed the presence of supernumerary Orb-positive cells in about 30% of *Hecw^CI^* mutants (**Fig. 3b**), as well as discrete Orb-positive puncta in the cytoplasm of nurse cells in both *Hecw^CI^* and *Hecw^KO^* mutant flies (**Fig. 4a**). To characterise the origin of Orb-positive puncta, we generated *Hecw^KO^* mutant flies expressing the RNP marker Me31B::GFP (ref. ^22^). In wild-type flies association of Orb with RNPs selectively occurs in the oocyte^23,24^. By contrast, in *Hecw* mutant animals altered Orb puncta fully colocalise with Me31B in the cytoplasm of nurse cells of previtellogenic and early vitellogenic egg chambers (**Fig. 4b**). Importantly, *Hecw^KO^* egg chambers contain significantly larger Me31B::GFP RNPs when compared to control (**Fig. 4b,d, Supplementary Fig. 5a**). Enlarged Me31B-positive RNPs are visible also in neurons of *Hecw^KO^* young adult brain (**Fig. 4c**).

**Figure 4.**
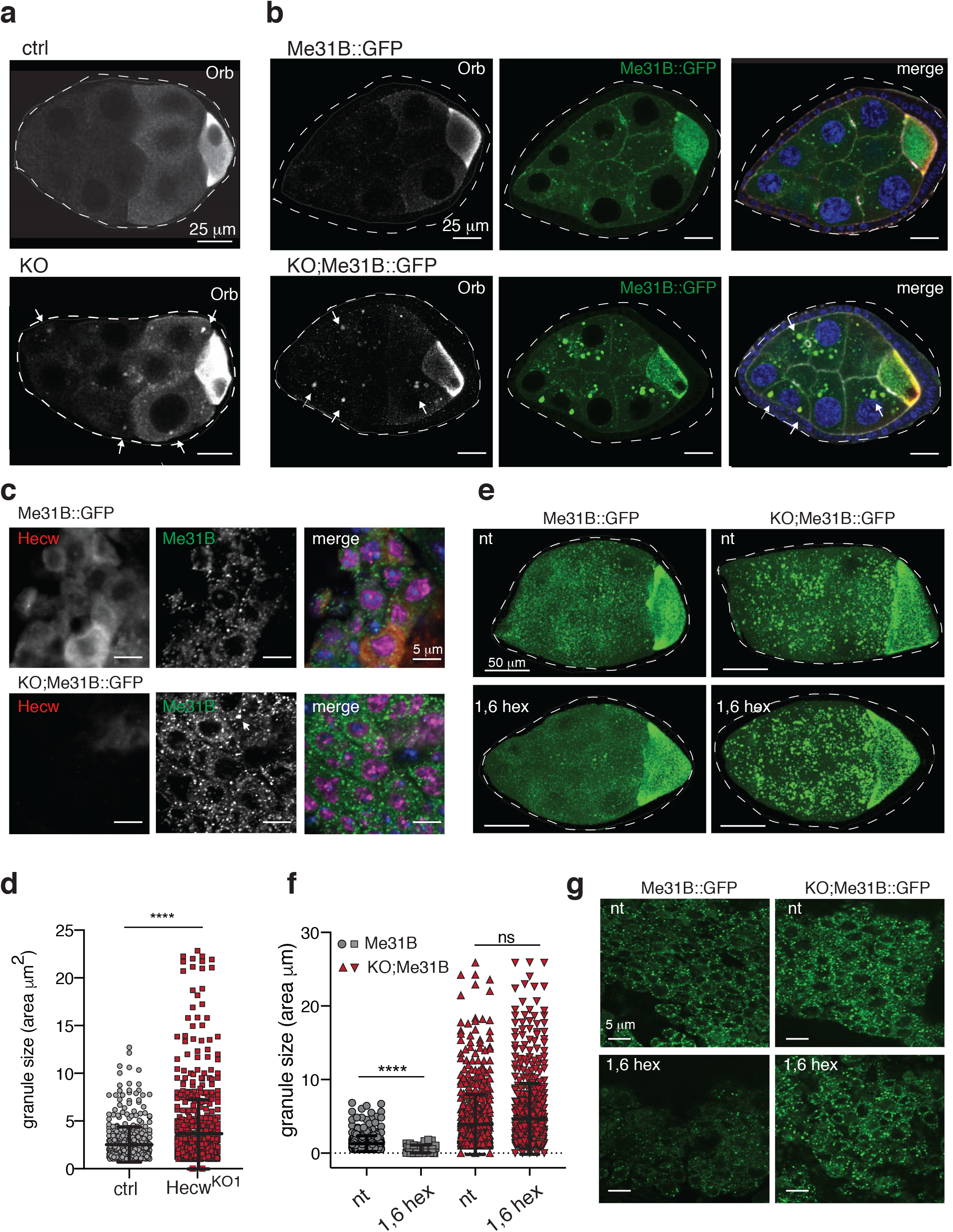
RNP morphology and composition is altered in Hecw^KO^. (**a**) IF analysis of egg chambers from 3-day-old flies of the indicated genotypes with anti-Orb antibody. White arrows indicate ectopic Orb puncta in nurse cells of *Hecw^KO^* flies. (**b**) IF analysis, as in a. Green, Me31B::GFP which marks the RNPs; red, Orb which marks the oocyte (in white in the first panel); blue, DAPI. Scale bar: 25 μm. (**c**) IF analysis of adult brains from 1-day-old flies with the indicated antibodies. Neurons are marked with the anti-elav antibody (purple). red, Hecw; blue, DAPI. Scale bar, 5 μm. (**d**) Quantification of RNPs size of egg chambers from 3-day-old flies of the indicated genotypes. Results are expressed as mean ± SD. *n*=477 (ctrl Me31B::GFP), *n*=559 (Hecw^KO^;;Me31B::GFP). 5 egg chambers/genotype, stages from 6 to 9 from 2 biological replicates. **** P<0,0001 by Mann-Whitney test. (**e**) IF analysis of egg chambers of 3-day-old flies as indicated, not treated (nt) or treated with 1,6 hexanediol (1,6 hex). Scale bar: 50 μm. (**f**) Quantification of RNPs size in egg chambers (stage 7 to 9) of the indicated genotypes and conditions. Treatment significantly reduces RNP size in control but not in *Hecw^KO^* flies (KO;Me31B::GFP). Particles analysed: *n*=401 ctrl (nt), *n*=38 ctrl (1,6 hex), *n*=642 *Hecw^KO^* (nt), *n*=553 *Hecw^KO^* (1,6 hex) (3 egg chambers/genotype). Size mean ± SD is reported. ****P<0.0001 by Mann-Whitney test. (**g**) IF analysis of adult brains from 1-day-old flies of the indicated genotypes, not treated (nt) or treated with 1,6 hexanediol (1,6 hex). Scale bar: 5 μm.

The dynamic composition of germline RNPs is linked to the biophysical status of membraneless organelles, which allows free diffusion in and out of the particles^25–27^. To investigate the nature of RNPs dynamics in the presence or absence of Hecw during *Drosophila* oogenesis, we performed live imaging of *Me31B::GFP* or *Hecw^KO^;Me31B::GFP* egg chambers (**movie 1,2**). Wild-type RNP particles appear mostly spherical and occasionally undergo fusion (**movie 3** and **Supplementary Fig. 5b**), two features of organelles with liquid-like properties^28,29^. This behaviour is altered in *Hecw^KO^* RNPs, which grow more irregularly shaped aggregates during the coarsening process due the *ex-vivo* observation **(movie 2** and **Supplementary Fig. 5c**).

To determine the nature of the interactions within RNP particles, egg chambers were treated with 2,5% 1,6 hexanediol, an aliphatic alcohol that disrupts weak hydrophobic bonds, typical promoters of liquid droplets^30–32^. While the size and number of wild-type granules is drastically reduced upon alcohol treatment, the larger *Hecw^KO^* RNPs barely dissolve, mirroring their solid-like nature (**Fig. 4e,f**). Similar results were obtained in neurons (**Fig. 4g**).

This result prompted us to evaluate the movement of Me31B particles, which is a combination of free diffusion and active, microtubule-dependent transport^33^. Live observation revealed no statistical differences in overall granule movement between control and *Hecw^KO^* flies. RNP mobility was quantified with particle tracking and with Differential Dynamic Microscopy (DDM) analysis^34^ to obtain the instantaneous velocity, the average mean square displacement *MSD*(Δ*t*) and the effective diffusion coefficient of RNPs. Results indicate that RNP transport is not impaired in the absence of Hecw (**Movie 1,2** and **Supplementary Fig. 5d-g**). Consistently, immunofluorescence analysis of the microtubule cytoskeleton structure in *Hecw^KO^* egg chambers showed no major alteration when compared to control (**Supplementary Fig. 6a**). Fluorescence recovery after photobleaching (FRAP) experiments confirmed that the dynamics of Me31B cytoplasmic fraction are not altered (**Supplementary Fig. 6b,c**).

To measure the exchange rate of Me31B::GFP molecules in RNP particles, we performed fluorescence loss in photobleaching (FLIP) measurements. The mobile fraction of Me31B::GFP is clearly reduced in *Hecw^KO^* egg chambers (**Fig. 5a**). Examination of single particles showed a significant difference in fluorescence decay between wild-type and *Hecw^KO^* flies (τ_WT_=(1.3±0.1)∙10^2^*s* versus τ_KO_=(4.2±0.1)∙10^2^s. P<10^−6^, **Fig. 5b**). Since we matched control and *Hecw^KO^* egg chambers for size or spatial distribution of RNPs (**Supplementary Fig. 6d-f**), this behaviour could only be attributed to a reduction in the exchange rate of the fluorescent protein between the RNP interior and the cytoplasm. This alteration is limited to the granules as the movement of cytoplasmic Me31B::GFP is comparable between wild-type and *Hecw^KO^* flies (**Fig. 5c**). Thus, in the absence of Hecw, RNP particles may undergo a transition to a less fluid, gel-like state.

**Figure 5.**
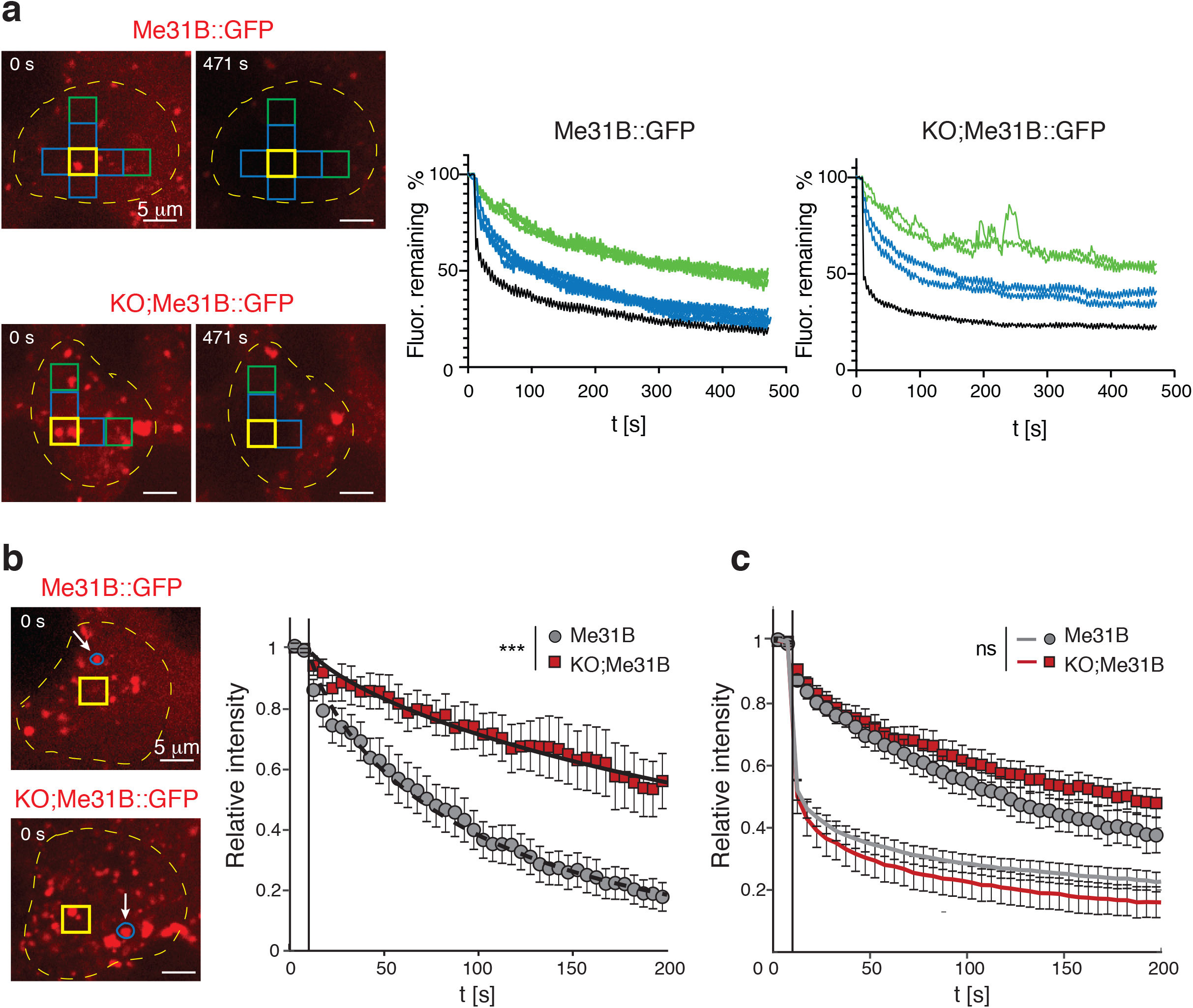
Lack of Hecw alters RNP biophysical proprieties. (**a**) FLIP analysis performed in *Me31B::GFP* or *Hecw^KO^ Me31B::GFP* (KO;Me31B::GFP) fly ovaries. Yellow boxes indicate the regions continuously bleached for 7 minutes, blue and green boxes the neighbouring regions in which Me31B::GFP fluorescence is recorded every 2 seconds. Right panel: example of fluorescence fluctuation in neighbouring color-coded regions, plotted as percent relative to time 0. The black line corresponds to the bleached ROI. *n*=11 egg chambers/genotype. (**b,c**) FLIP analysis performed as in a, by quantifying the fluorescence of single RNPs (**b**, white arrow in the example) present in neighbouring regions. Relative intensities of Me31B::GFP fluorescence in RNPs are plotted relative to time. *n*=18 RNPs/genotype. Data are reported as mean ± SD. ****P<0.0001 by Mann-Whitney test. (**c**) Upper curves: FLIP curves in the cytoplasm. Lower curves: relative intensity within the bleached area. Both in the cytoplasm and in the bleached area, the loss curves show no statistically significant difference (P =0.07 by t-student) between control and *Hecw^KO^*.

Overall, these results indicate that Me31B::GFP-positive RNPs possess liquid droplet properties that are regulated by Hecw-mediated ubiquitination.

### Hecw interacts and colocalises with RNPs components

The altered physical properties of Me31B::GFP RNPs prompted us to investigate whether Hecw may interact with RNP proteins. Consistently, in ovarian extract, Hecw co-immunoprecipitates with Me31B::GFP together with other known RNP components, such as Orb and Fmrp (ref.^5^) (**Supplementary Fig. 7a**). The interaction is maintained upon RNAse treatment, suggesting that Hecw interacts directly with RNPs via protein-protein interactions (**Fig. 6a**).

**Figure 6.**
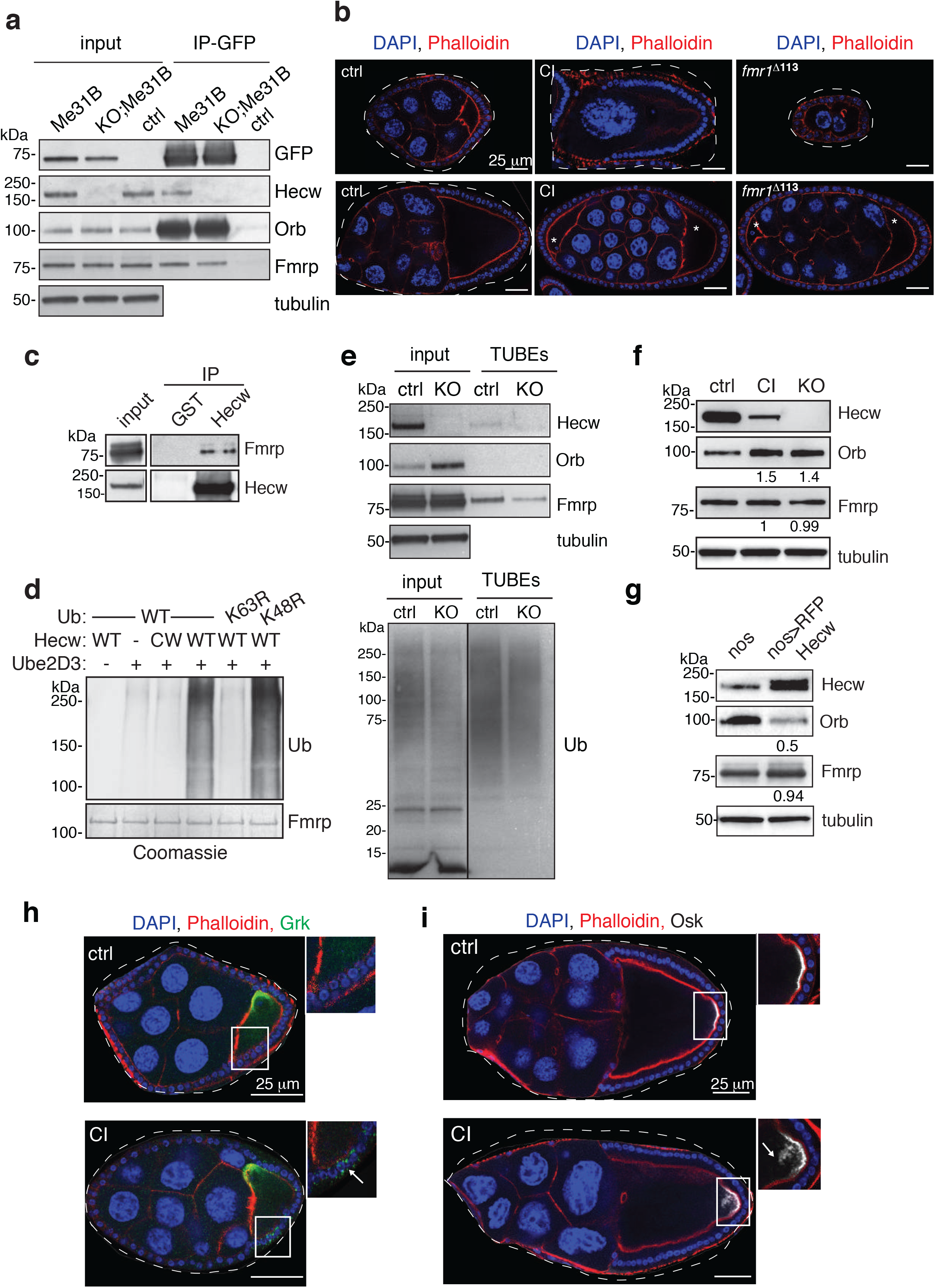
Hecw interacts with RNP components and ubiquitinates Fmrp. (**a**) 0,5 mg ovary lysates from *Me31B::GFP* (Me31B), *Hecw^KO^1;Me31B::GFP* (KO;Me31B) and control (yw) lines were IP with anti-GFP antibody in the presence of 100 ug/ml RNAseA. Input correspond to 2% of the total immunoprecipitated proteins. IB as indicated. (**b**) IF analysis of egg chambers from 3-day-old wild-type (ctrl, left), *Hecw^CI^* (CI, middle) or *Fmr1* mutant (*Fmr1*Δ113, right) flies. Mutant flies present same defects with similar penetrance. Examples of reduced number of nurse cells (upper panels) and compound egg chambers (bottom panels) are reported. Red, phalloidin; blue, DAPI. Asterisks indicate oocytes of the compound egg chamber. Scale bar: 25 μm. (**c**) 0.5 mg of ovary lysates from the indicated lines were IP and IB as indicated. Input corresponds to 1% of total immunoprecipitated proteins. (**d**) *In vitro* ubiquitination assay of Fmrp with ubiquitin wild-type, K63R or K48R mutants analysed by IB with anti-Ub antibody. Coomassie shows equal loading. (**e**) 0.6 mg of ovary lysates from the indicated lines were IP for Ub with TUBEs and IB as indicated. Input correspond to 3% of total immunoprecipitated proteins. (**f**) IB of wild-type (ctrl)*, Hecw^CI^* (CI) and *Hecw^KO^* (KO) fly ovaries with the indicated antibodies. Orb levels are increased upon Hecw depletion. Values reported above each blot are normalised for tubulin and represent the fold change relative to the control, presented as mean of three experiments. (**g**) IB of control (driver only, *nos-GAL4*, nos) and RFP-Hecw-overexpressing fly ovaries (*nos>RFP Hecw*) with the indicated antibodies. Orb levels are decreased upon Hecw overexpression in the germline. (**h,i**) IF analysis of egg chambers of 3-day-old wild-type and CI with the indicated antibodies. White arrows in the magnifications indicate mislocalised Grk (green, **h**) at the ventral side of the oocyte or mislocalised Osk (white, **i**) diffused from the posterior margin of the oocyte in the Hecw mutant flies. Scale bar: 25 μm.

To identify Hecw binding partners, we performed a pull-down assay using a GST construct spanning the two WW domains as a bait, followed by mass spectrometry (**Supplementary Table 2**). The majority of Hecw interactors are mRNA binding proteins involved in multiple steps of RNA processing (**Supplementary Table 2**), among them the validated Fmrp. Strikingly, other two interactors, Hrp48 and Glorund, are translational repressors previously implicated in translational control during oogenesis^35^. These findings provide compelling evidence for a critical role of Hecw in RNPs regulation.

### Hecw-dependent ubiquitination modulates Orb via Fmrp

How could Hecw-dependent ubiquitination regulate RNPs? We focused on Fmrp, a known RNA binding repressor of the Orb autoregulatory loop in nurse cell^20^. *Fmr1^Δ113^* loss of function mutant flies^20^ display oogenesis defects that resemble the ones of *Hecw* mutants (**Fig. 6b**). Corroborating the idea that the two proteins act on the same axis, genetic interaction revealed no worsening of the phenotypes (**Supplementary Fig. 7b**). Additionally, we found that endogenous Hecw and Fmrp from fly ovaries coimmunoprecipitate (**Fig. 6c**) and that Fmrp is a substrate of Hecw, as measured by *in vitro* ubiquitination assay (**Fig. 6d**). We confirmed this result *in vivo* by isolating endogenous ubiquitinated targets using tandem-repeated Ub-binding entities (TUBEs), a sensitive method that demonstrates the covalent attachment of polyubiquitin chains to proteins modified at low stoichiometry^36^. While Fmrp is clearly present in the pull-down from wild-type ovaries, the lane corresponding to the *Hecw^KO^* ovaries displays a remarkably reduced amount of protein (**Fig. 6e**), indicative of reduced Fmrp ubiquitination in the absence of the E3 ligase.

To evaluate the impact of Hecw-dependent ubiquitination on Fmrp, we examined its protein levels in wild-type and *Hecw* mutant ovaries. As predicted by Hecw specificity towards K63-linked polyubiquitination (**Fig. 6d**), Fmrp stability is not affected by the lack of Hecw activity (**Fig. 6f**). In contrast, the downstream target Orb accumulates in *Hecw* mutant ovaries (**Fig. 6f**), in good agreement with ectopic presence of Orb puncta in nurse cells (**Fig. 4a,b**). Regulation by Hecw appears to occur at the protein level, since both *Fmr1* and *orb* mRNA expression does not change significantly when compared with wild-type controls (**Supplementary Fig. 7c**). Consistent with the antagonistic effect exerted by Hecw on Orb through Fmrp, overexpression of Hecw in the germline strongly downmodulates Orb protein levels (**Fig. 6g**) without affecting its mRNA (**Supplementary Fig. 7d**) and treatment of ovaries with the proteasome inhibitor MG132 does not rescue Orb expression (**Supplementary Fig. 7e**).

Me31B-positive RNPs in ovaries control spatial and temporal translation of key mRNAs during development^22^. To test whether the aberrant RNP structure and composition of *Hecw* mutant tissue alters translation control, we analysed the localisation of the two major Orb target proteins: Gurken (Grk) and Oskar (Osk). While the mRNA levels are not altered (**Supplementary Fig. 7c**), both proteins show aberrant localisation in *Hecw* mutant oocytes. Grk is found ectopically in the ventral portion of the oocyte (**Fig. 6h**), while Osk localises away from the posterior margin (**Fig. 6i**). Remarkably, in stage 8/9, ectopic Osk colocalises with Me31B::GFP positive particles in *Hecw* mutant egg chambers, suggesting premature translation (**Supplementary Fig. 7f**), as is the case of Orb. Overall, these data indicate that Hecw is a positive regulator of Fmrp repressor activity on Orb and its targets.

## DISCUSSION

We report here the identification and molecular characterisation of Hecw, the fourth member of the NEDD4 family in *Drosophila*. Flies lacking Hecw activity show neurodegenerative traits, shorter lifespan and reduced fertility. We provide compelling evidence that Hecw-mediated ubiquitination is needed for the diffusive exchange in and out of RNP compartments during oogenesis and in neurons. Mechanistically, Fmrp ubiquitination by Hecw is required for the translational repression of Orb, a Fmrp target and oocyte determinant.

### Ubiquitin controls RNPs liquification

RNP granules play a fundamental role in both germ and somatic cells as they are responsible for the mRNA transport and local translation required for neuronal and oocyte maturation^5,37,38^. Among the various RNPs, sponge bodies are marked by the de-capping activator/RNA helicase Me31B (ref.^22^). These are first synthesised in nurse cells and then transported and localised posteriorly in developing oocytes, where their mRNAs are translated, dictating the establishment of germ cells and embryonic axes^39–41^. While their composition has been recently investigated^42,43^, the mechanism behind the dynamic nature and biophysical properties of sponge bodies are largely unknown. Recent literature underlined a liquid-like behaviour for RNP granules in *Caenorhabditis elegans*^25,44^, as well as for germ granules in fly embryos^45^. We now show that also *Drosophila* sponge bodies behave like liquid droplets.

Our data provide evidence for the pivotal role of Hecw and ubiquitin in maintaining the liquid-like nature of RNPs, and show that, in the absence of Hecw, they transition into a less dynamic gel-like state. The lower exchange rate between RNPs and the cytoplasm is specific of *Hecw* mutant granules and possibly reflects an altered internal rearrangement and activity of the proteins present inside these membrane-less particles. Indeed, the Orb protein that is usually absent in the nurse cells granules^23,24^, is clearly present in the larger Hecw-depleted RNPs, suggesting a lack of local translational repression. Although our study is limited to the analysis of Me31B in oocytes and neurons, it is conceivable that other RNPs behave similarly and that ubiquitination may modulate liquification of other phase-separating complexes, promoting specific regulated events. This seems to apply to UBQLN2-dependent stress granules, in which ubiquitin appears to enable shuttling of client proteins out of the granules^46^. It is important to note that, as for other ubiquitin-modulated pathways^47^ specific types of ubiquitination (e.g. K63- or K48-linked chains) may easily drive different outcomes, ultimately adding another exciting layer of complexity to explore in the future.

### Hecw acts in the Fmrp-Orb axis

We identify Fmrp as interactor, substrate and effector of Hecw activity in RNPs. Importantly, a Hecw mutant phenocopies Fmrp loss of function flies, both in terms of phenotypic defects and molecular target. Indeed, Orb, a known target of Fmrp repressor activity^20^, shows an inverse correlation with Hecw expression, an effect that cannot be ascribed to a direct activity of Hecw on Orb. Similar to Fmrp (ref.^48^), changes in Hecw expression levels are deleterious and induce pathological effects. Based on this evidence, we propose a linear cascade of events in which Hecw ubiquitinates Fmrp to maintain mRNAs of Orb in a repressed state. Such K63-linked, non-proteasomal function of Hecw in Fmrp ubiquitination is in good accordance with the previously reported activity of this enzyme family^8,9^, and is clearly different from the proteasome-mediated degradation of Fmrp identified in neurons^49^.

Fmrp represses translation both during ovary and neuronal development, however, the underlying mechanism remains unclear^50,51^. We envisage that ubiquitin, by adding an additional surface of interaction to Fmrp, could enable precise spatiotemporal recruitment of repressive components^52^ and ribosomes^53^. Indeed, while *Drosophila* Fmrp directly binds the 80S ribosome near the tRNA and translation factor binding site, post-translational modifications (PTMs) have been suggested to modulate the affinity of Fmrp for the ribosome or target mRNAs, thereby ‘‘turning off’’ protein synthesis locally^53^. Thus, a ubiquitin-dependent inhibition of Fmrp association to ribosomes might occur in nurse cells and be released by deubiquitination in the oocyte when translation is due to progress. Similar cycles of ubiquitination/deubiquitination have been identified and characterised in other cellular processes ^54,55,56^. Thus, the existence of Ub receptors, effector components capable of recognising ubiquitin, is predicted^57^. One such receptor may well be the P-body component Lingerer, a protein endowed with a ubiquitin associated (UBA) domain known to associate with Fmrp and other RNA-binding proteins to modulate their repressive functions^58^, which we identified as a Hecw interactor (**Supplementary Table S2**).

While our data are fully compatible with this model, it is also conceivable that Ub may impinges on the physical properties of Fmrp inside RNP granules. Recent studies showed that phosphorylation, methylation and sumoylation could control Fmrp phase-separation propensity, and regulation of its intrinsically disordered regions^59–62^. In the less mobile *Hecw* mutant RNPs, Fmrp may be disordered or otherwise uncapable of interacting with repressive complexes and ribosomes. Further studies are needed to test the behavior of ubiquitinated vs non-ubiquitinated Fmrp and define functional links with different PTMs.

### Hecw and disease

*Hecw* mutant flies show premature aging, neuronal loss and neuromotor defects, all reminiscent of neurodegenerative processes in humans. The biophysical properties and dynamics of RNPs in neurons are predicted to be affected by the absence of Hecw, which may severely impact on neuronal homeostasis. Indeed, mutations in critical RNP components^63,64^ have all been extensively linked to neurological diseases such as amyotrophic lateral sclerosis (ALS) and frontotemporal dementia (FTD), highlighting the crucial contributions of RNA localisation and translational control to long-term neuronal integrity^65,66^. Furthermore, liquid-to-solid–state transitions are considered the key event in the formation of intracellular pathological protein aggregates^63,67^.

Remarkably, scattered evidences links the human ortholog HECW1 to neurodegeneration. Transgenic mice overexpressing HECW1 show loss of neurons in the spinal cord, muscular atrophy and microglia activation^68^, and HECW1 protein has been shown to ubiquitinate mutant superoxide dismutase 1 (SOD1), typical of familial ALS patients^10^. Similar to Hecw, the expression of HECW1 seems to decline with aging^69^, suggesting the existence of a positive, protective role for this E3 ligase in the maintenance of neuronal homeostasis.

In humans, FMR1 alterations are at the basis of the Fragile X syndrome, characterised by intellectual disability, autism as well as premature ovarian failure^70^. Abnormally expanded CGG segment in the promoter silences at various degree the *FMR1* gene located on chromosome X, causing a spectrum of pathological phenotypes. Based on our findings, we can hypothesise that the disorder severity may also depends on Hecw status both in neurons and in ovaries. Future studies on Hecw/HECW1 in flies and mammalian models are predicted to illuminate new pathogenetic aspects of this and other diseases.

## METHODS

### *Drosophila* strains and cell lines

Flies were maintained on standard flyfood containing cornmeal, molasses and yeast. All experiments were performed at 25°C, unless differently specified. The following fly strains were used in this study*: y^1^w^1^* (Bloomington *Drosophila* Stock Center [BDSC] #1495), *nanosGAL4-VP16* (kindly provided by A. Ephrussi), *elav-GAL4* (BDSC #25750), *traffic jam-GAL4* (Kyoto stock center, [DGRC]#104055), *Me31B::GFP* (ref.^71^, gift of Tim Weil), *Fmr^1Δ113M^* ([BDSC] # 6929), *UAS-Hecw^RNAi^* CG42797 RNAi (Vienna *Drosophila* Stock Centre [VDRC] #104394). The rescue line *Hecw^KO;;DC504^* was generated by crossing *Hecw* mutant flies with the *Dp(1;3)DC504* (BDSC #32313), which contain a duplication of *CG42797* on the third chromosome. Transgenic flies were generated by injecting the UAStattB-RFP-CG42797 construct into embryos carrying the attP-zh86Fb φC31 docking site (BestGene.inc). Transgenic offspring was screened by eye-colour (white marker) and sequenced. Strain details are reported in **Supplementary Table 3**. *Drosophila* S2 cells obtained from Invitrogen, were cultured in Schneider’s medium (GibCO) supplemented with 1% Glutamine (Euroclone) and 10% of Fetal Bovine Serum (FBS) (Euroclone) and maintained at standard culture conditions (28 °C).

### *Hecw^CI^* and *Hecw^KO^* and generation by CRISPR/Cas9 editing

Guide sgRNAs for CRIPR/Cas9 mutagenesis were designed using the MIT CRISPR design tool (http://crispr.mit.edu, Zhang Lab, MIT). Target sequences were cloned in the pBFvU6.2 vector (NIG-fly stock center) between two BbsI restriction sites with the following oligos:

Hecw^CI^ 5’-<u>CTTC</u>GTGCCCACACATGCTTCAAT-3’, 5’-<u>AAAC</u>AATGAAGCATGTGTGGGCAC-3’.
Hecw^KO^ 5’-<u>CTTC</u>GCCTTCTACGAGGCGCGCAA-3’, 5’-<u>AAAC</u>TTGCGCGCCTCGTAGAAGGC-3’.

The sgRNA constructs were injected into *y,w P{nos-phiC31}; attp2* embryos. Transformants sgRNA flies were crossed with *y^2^ cho^2^ v^1^; P{nos-Cas9, y+, v+}3A/TM6C, Sb Tb* (DGRC # CAS-0003) to obtain founder animals with both transgenes. Founder males were crossed with compound-X chromosome (BDSC #64) and the potentially mutated chromosomes were recovered from founder animals over FM7. Cas9 and sgRNA elements were removed from the background thanks to selection of v+ eye colour. Mutated chromosomes were identified using T7EI assay. Mutations were further characterised by PCR amplification and sequencing of the target region with the following primers:

Hecw^CI^ F: 5’-CCGAGAGTTGGAGCTGGTTA-3’, R: 5’-AAACTAGTGGGATGCCATGC-3’.
Hecw^KO^ F: 5’ ATGGAGCCACCAGCT 3’, R: 5’ AGCTGGTGGCTCCAT 3’.

For alleles details see **Supplementary Fig. 2**.

### Constructs

To generate the pUAStattB-RFP-Hecw construct, the *Hecw* gene was amplified from LD10978 vector (*Drosophila* Genetic Resource Consortium [DGRC]), with the following primers: CG42797BglII_F: 5’-GTCCGGACTCagatctATGGAGCCACCAGCTGCA-3’ and CG42797XhoI_R: 5’-TAGAGGTACCctcgagCTACTCAATGCCGAACGTGTTG-3’ and cloned by enzymatic digestion into a pUAStattb/RFP vector, previously created using the Infusion HD cloning system (Takara Clontech).

To generate the pGEX6P1-Hecw wild-type construct, the full-length Hecw was amplified from pUAStattB-RFP-Hecw with the following primers: CG42797EcoRI_F: 5’-CCGgaattcATGGAGCCACCAGCT-3’ and CG42797XhoI_R: 5’-TAGAGGTACCctcgagCTACTCAATGCCGAACGTGTTG-3’ and cloned by enzymatic digestion into a pGEX6P1(GE Healthcare).

The pGEX6P1-Hecw C1394W construct was generated by site-directed mutagenesis according to the QuikChange Site-Directed Mutagenesis Kit protocol (Agilent) using the following primers: F: 5’-CCCGTGCCCACACATGGTTCAATCGGCTGGATTTG-3’ and R: 5’-CAAATCCAGCCGATTGAACCATGTGTGGGCACGGG-3’.

To generate pGEX6P1-WW construct, the region containing the two WW domains of Hecw (636-834aa) was amplified from pUAStattB-RFP-CG42797 construct with the following primers: CG42797EcoRI WWI_F: 5’-ccgGAATTCCCACCATTGCCGCCTG-3’ CG42797XhoI WWII_R: 5’-ccgCTCGAGTCAACGAGGATCCATGAA-3’ and cloned by enzymatic digestion into a pGEX6P1(GE Healthcare).

To generate a pGEX6P1-Hecw (1-130aa) construct for use in antibody production, the N-terminal region of the protein was amplified from pUAStattB-RFP-CG42797 with the following primers: CG42797EcoRI_F: 5’-CCGgaattcATGGAGCCACCAGCT-3’ and CG42797XhoI_Nterm_R: 5’-CCGctcgagTCATTCGCTGGGCTGC-3’ and cloned by enzymatic digestion into a pGEX6P1(GE Healthcare).

To generate pET43-His-MBP-Fmrp construct, full-length Fmrp was amplified from cDNA of *y^1^w^1^* ovaries using the following primers: Fmrp BamH_F: 5’-CGCggatccATGGAAGATCTCCTC-3’ and Fmrp EcoR1_R: 5’-CCGgaattcTTAGGACGTGCCATT-3’ and cloned by enzymatic digestion into a pET43-His-MBP vector (kind gift of Sebastiano Pasqualato, European Institute of Oncology, Milan).

All constructs were sequence-verified. The remaining constructs were previously described ^9^.

### RNA extraction and qPCR

RNA was extracted from *Drosophila* tissues with Maxwell RSC simplyRNA Tissue kit (Promega) and quantitative PCR (qPCR) analysis was performed with the following TaqMan Gene Expression Assay (Thermo Fisher Scientific): Dm01837441_g1 (*CG42797/Hecw*), Dm01841193_g1(*gurken*), Dm02136373_m1(*Fmr1*), Dm02136342_g1(*orb*), Dm02134538_g1 (oskar).

### Lifespan Assay

1 to 3-day-old flies (100 males and 100 females) were kept at 25°C or 29°C at a density of 25 flies/vial, in mixed-sex groups. Flies were flipped and scored every two/three days for survivorship. The assay was repeated twice, and data were analysed with PRISM (GraphPad software). Survival fractions were calculated with product limit Kaplan-Meier method and log rank test was used to evaluate the significance of differences between survivorship curves.

### Climbing Assay

8 flies/genotype were placed in a 9 cm plastic cylinder. After a 30-seconds rest period, flies were tapped to the bottom of the cylinder. Negative geotaxis was quantitated by counting the number of flies that can cross a 7 cm threshold during a 15-seconds test period. The climbing index was calculating as the number of succeeding flies over the total. The test was performed on 8 groups of animals for each genotype. The climbing ability was measured the first day of life and monitored every 5 days. Significant differences between *Hecw* mutants and control started to emerge after the 14^th^ day of the lifespan.

### Fertility assay

Fertility was assessed by counting the eggs laid in 24 hours by 20-day-old flies. 6 females and 3 males per genotype were kept in a cage on a 3 cm molasses plate with fresh yeast for a 24-hour egg collection after 24 hours of adjustment to the cage. The test was performed on 3 groups/genotype. The hatching rate was calculated as the number of eggs hatched into larvae in 3 day over the total number of eggs.

### Immunostaining and treatments

Fly ovary dissection and staining was performed as previously described ^72^. Briefly, ovaries were fixed in 4% paraformaldehyde (PFA) in PBS (HIMEDIA) for 20 minutes at room temperature, permeabilised in 1% Triton X-100 in PBS for 20 minutes, followed by 1-hour block in PBST (PBS-Triton X 0.1%) containing 5% (w/v) bovine serum albumin (BSA). Primary antibodies were incubated overnight at 4°C, secondary antibodies were incubated for 2 hours and DAPI (Sigma) was incubated for 15 minutes at room temperature. Adult brains (**Fig. 1e, Fig. 4d**) were dissected in *Drosophila* S2 medium with 10 % of Fetal Bovine Serum (FBS) (Euroclone) and processed as described before. Ovary microtubule detection (**Supplementary Fig. 6a**) was adapted from^73^. Briefly, ovaries were incubated in BRB80-T buffer (80 mmol PIPES pH 6.8, 1 mmol MgCl2, 1 mmol EGTA, 1% Triton X100) for 1 hour at 25°C without agitation. Then, ovaries were fixed in MeOH at –20°C for 15 minutes and rehydrated for 15 hours at 4°C in PBST and blocked for 1 hour in PBST containing 2% (w/v) BSA before incubation with primary and secondary antibodies overnight in PBST-2% BSA.

In case of drug treatments, fly ovaries were dissected in *Drosophila* S2 medium with 10 % of FBS (Euroclone) and incubated with 2.5% 1,6-hexanediol on an orbital shaker at room temperature for 5 minutes (**Fig. 4e**) prior to fixation.

All tissues were mounted in 20% glycerol, 50 mM Tris, pH 8.4 to avoid mechanical sample deformation. Images were acquired using 40X and 60X objectives on a Leica TCS Sp2 upright confocal microscope. The outlines of the oocytes and the anteroposterior (AP) axis were manually specified.

For S2 immunostaining (**Supplementary Fig. 1e**), cells were plated on coverslips coated with poly-ornithine. S2 cells were rinsed twice with PBS and fixed in 4% paraformaldehyde (PFA) for 15 minutes, permeabilised with PBST for 20 minutes. After 30 minutes incubation in Blocking solution (PBST-1% BSA), coverslips were incubated with primary antibody diluted in PBST 0,1% BSA for two hours at room temperature. After three washes in PBS, cells were incubated with secondary antibodies for 2 hours at room temperature. DAPI (Sigma) was incubated for 10 minutes at room temperature then washed once in PBS. Coverslips were mounted on slides using Mowiol Mounting Medium (Calbiochem).

TUNEL staining was performed as previously described^74^. Briefly, fixed brains from 30-day-old flies were permeabilized in 100 mM sodium citrate, 0.3% Triton X-100 PBS at 65°C for 45 minutes. Brains were then incubated with TUNEL reagent (In Situ Cell Death Detection Kit, TMR red, Sigma) for 14-16 hours at 37°C in dark humid chamber, washed in PBST and incubated with HOECHST 33342 (Life Technologies, 2μg/ml in PBS) 10 minutes at 25°C. Finally, brains were rinsed in water and mounted on glass slides with Prolog Gold fluorescence anti-fading reagent (Invitrogen). Images were acquired with a Nikon ECLIPSE C1si confocal microscope. To quantify TUNEL+ cells, 200-500 cells/individual from *n*=5 individuals/genotype were scored using the FIJI Software (https://imagej.net/Fiji).

### Immunohistochemical analysis

Adult heads of 1-day, 30-day and 60-day-old flies were dissected in PBS and fixed in 4% PFA (HIMEDIA) overnight, at 4°C. Samples were embedded in 1,2% low-melting agarose: while the agarose solidified, heads were properly oriented. Heads-containing agarose blocks were dehydrated in serial dilutions of ethanol (from 70% to 100%) prior to paraffin embedding with a Leica ASP300 Enclosed Tissue Processor. The paraffin blocks were cut with a Leica RM2125 RTS Microtome into 5 μm frontal sections, stained with haematoxylin–eosin (HE) according to standard procedures, and examined by bright-field microscopy. For each time point, at least 5 brains/genotype were analysed, and vacuoles with diameter >2 μm were counted over 12/15 brain slices.

### Live imaging of GFP labelled protein

For *ex vivo* live recording, ovaries of 3-day old *Me31B::GFP* and *Hecw^KO^;Me31B::GFP* females were dissected and mounted on glass bottom MatTek (35mm) in Halocarbon oil 27 (Sigma) according to ^75^. Live cell imaging experiments were performed on the UltraVIEW VoX spinning-disk confocal system (PerkinElmer), equipped with an EclipseTi inverted microscope (Nikon), a Hamamatsu CCD camera (C9100-50) and driven by Volocity software (Improvision; Perkin Elmer), using a 40x oil-immersion objective (Nikon Plan Fluor, NA 1.3) and a 488 nm laser. One frame every 10 seconds was acquired for a maximum of 30 minutes to avoid phenotypic changes due to stress.

### RNP velocity, size and shape measurements

Me31B::GFP RNP velocity and size analysis was performed using Fiji^76,77^. Particles were tracked with the Manual tracking plugin of Image J and analysed with the Chemotaxis plugin (https://ibidi.com/chemotaxis-analysis/171-chemotaxis-and-migration-tool.html).The cursor was placed at the leading edge of each particle in the direction of the movement and tracked until the particle moved out of the plane of focus or ceased moving. The X and Y position of each particle was recorded. Particle size was measured using the Analyze particle tool, setting the same threshold for all images.

The circularity of the RNPs was quantified by evaluating for each granule the so-called shape factor, which is defined as 4*πA*/*P*^2^, where *A* is the projected area of the granule and *P* its perimeter. The definition is such that the shape factor takes the value 1 for a perfectly round object, while it assumes smaller and smaller values as the object become more elongated or irregularly shaped. Area and perimeter of each RNP in a given ovary were obtained by processing the corresponding confocal image *via* a custom MATLAB® code using the function bwboundaries (with 4 pixels connectivity), applied to a binary image obtained from the original image by thresholding. In the case of time-lapse acquisitions, the same threshold was used for all the images in the sequence.

### Fluorescent Recovery After Photobleaching (FRAP)

For FRAP recordings, 3-day old *Me31B::GFP* and *Hecw^KO^;Me31B::GFP* females were prepared as described above. Time series were recorded using a Confocal Spinning Disk microscope (Olympus) equipped with IX83 inverted microscope provided with an IXON 897 Ultra camera (Andor) and a IX3 FRAP module equipped with a 405 nm laser using 60X objective. The system is driven by the Olympus CellSens Dimension 1.18 software (Build 16686). The recording frame rate of the GFP signal, excited by 488 nm laser, was 66 ms. Photobleaching was performed via the 405 nm laser for 2 cycles of 500 ms on a square region of the cytoplasm (3,04 x3,04 μm) to bleach GFP signal.

Before bleaching, 15 frames were acquired, obtaining a pre-bleaching reference image *I*_0_(***r***). After bleaching, time-lapse observation was continued for 1500 frames. We consider the azimuthally-averaged intensity profile of the bleached region 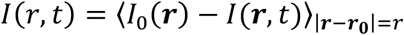, where *I*(***r***, *t*) is the image intensity at time t after bleaching, ***r***_0_ is the center of the bleached region and the symbol 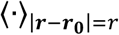 indicates an average over all pixels in the image at a distance *r* from the center ***r***_0_ of the bleached region. The process of fluorescence recovery was monitored by following the progressive widening of the bleached spot over time. For each fixed *t*, *I*(*r*, *t*) was fitted by a Gaussian profile *A*(*t*)*exp*[−*r*^2^/*σ*(*t*)^2^] + *B*(*t*), where the parameter *σ*(*t*) provides an estimate of the width of the bleached spot. In all cases, we found that the increase of over time of *σ*(*t*) was well captured by the simple equation 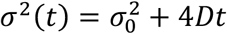, describing the diffusive-like broadening of a Gaussian concentration profile. From a linear fit of *σ*^2^(*t*) over a time interval of 30 seconds, an estimate of the effective diffusion coefficient *D* was obtained, as shown in the inset of **Supplementary Figure 6b**. Compared to standard data treatment, where the FRAP process is monitored by measuring the average intensity within the bleached region, the described procedure, which is based on the time-resolved estimation of a parameter obtained from the fit of a spatial feature, is more robust with respect to the large local intensity fluctuations in the images due to the randomly-moving, highly-contrasted RNPs, whose appearance in the vicinity of the bleached ROI can significantly alter the shape of the recovery curve. The distributions of the obtained values of *D* are shown in **Supplementary Figure 6c**, showing no statistically significant difference between the wild-type and *Hecw^KO^* nurse cells (*D*_*WT*_ = (0.24 ± 0.03)*μm*^2^*s*^−1^, *D*_*KO*_ = (0.24 ± 0.02)*μm*^2^*s*^−1^).

### Fluorescent Loss in Photobleaching (FLIP)

FLIP experiments were performed on the UltraVIEW VoX spinning-disk confocal system (PerkinElmer) equipped with an EclipseTi inverted microscope (Nikon), provided with an integrated FRAP PhotoKinesis unit (PerkinElmer) and a Hamamatsu CCD camera (C9100-50) and driven by Volocity software (Improvision; Perkin Elmer). Photobleaching was achieved on a square region of 4×4 μm by using the 488 nm laser at the maximum output to bleach the GFP signal every 5 seconds, for a total of 100 bleaching events. Initially, five images were acquired to determine the levels of pre-bleach fluorescence. Images were acquired through a 60X oil-immersion objective (Nikon Plan Apo VC, NA 1.4) every 2.5 seconds for 8 minutes.

In order to analyse the fluorescence fluctuations over time in FLIP experiments, two homemade Fiji plugins were developed; one to analyze fluorescence either in selected and manual tracked RNPs or in the cytoplasm and the other one to evaluate the fluorescence in the bleached area and in its neighborhood regions. During each acquisition, a background intensity *I*_*R*_(*t*) and a reference intensity *I*_*B*_(*t*) were measured. The amplitude *I*_*R*_(*t*) of the background noise was obtained as the average intensity within a small ROI in a corner of the image outside the egg chamber. *I*_*B*_(*t*) monitors potential systematic changes in the fluorescent emission not directly related with the FLIP experiment and was calculated as the average intensity within a small ROI inside the sample, far from the bleached region and in a different nurse cell. In all of the experiments, we found that both *I*_*R*_(*t*) and *I*_*B*_(*t*) were fairly constant over time, with no increasing or decreasing net trends. *I*_*R*_(*t*) showed very small fluctuations (with RMS amplitude below 5% of the mean value), while in some cases, the profile of *I*_*B*_(*t*) was perturbed by the presence of granules moving in and out the ROI. Thus, we adopted a constant value *I*_*B*_ for the background intensity (obtained as a time average of *I*_*B*_(*t*)) and we did not apply further corrections for systematic intensity drifts.

In each experiment, we selected N (6<N<8) ROIs of average area 2.5 *μm*^2^ outside of the bleached area (half of them in the cytoplasm, half of them in correspondence to RNPs). The area of a ROI associated with an RNP was kept constant over time, while its position was adjusted in order to follow the moving granule. For the n-th ROI, the average intensity *I*_*n*_(*t*) was measured. The relative intensity was calculated as: 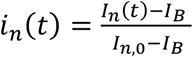, where *I*_*n*,0_ was the average intensity within the ROI before bleaching.

Since the diffused intensity can be larger in the region occupied by the cells compared to the outside of the egg chamber, our estimate of *I*_*B*_ represents a conservative estimate of the background contribution. In general, underestimating the background intensity can lead to a systematic overestimate of all relative intensity. However, we found that: i) the loss curves in the cytoplasm showed no statistically significant difference (in terms of amplitude and kinetics) between *Hecw^KO^* and wild-type; ii) the loss curves in the bleaching area showed no statistically significant difference (in terms of amplitude and kinetics) between *Hecw^K^* and wild-type; iii) the loss curves of the RNPs in *Hecw^KO^* and wild-type were significantly different, the decay time of wild-type RNPs being about 3 times faster, independently of particular choice of the background intensity. We explicitly verified this independence by repeating the analysis with different values of *I*_*B*_ (25% smaller than the originally estimated value, 25% larger, randomly chosen in in the interval [original value −25%, original value −25%]).

To extract a characteristic decay time *τ* from the fluorescence loss curves of the RNPs, we fitted our data to a single stretched-exponential decay 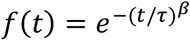, with *β* = 0.75.

A single laser exposure was not enough to completely “switch-off” the fluorescence signal within the photobleached ROI (Fig.5). In fact, the relative intensity drop was only about 0.5 for both *Hecw^KO^* and wild-type.

### Differential Dynamic Microscopy (DDM)

To quantify the mobility of RNPs, time series confocal images were analysed with DDM that enables the tracking-free characterisation of the dynamics in a variety of nano- and micro-systems where the individual objects cannot be resolved or tracked with sufficient accuracy, a typical scenario of biological samples^78^. Rather than reconstructing trajectories followed by the individual particles during their motion in direct space, DDM extracts quantitative mobility information from the analysis of the temporal correlations of the spatial Fourier transforms of the direct space images^79^. We used DDM to estimate the average mean square displacement *MSD*(Δ*t*) of the RNPs^34^.

For each image sequence, DDM analysis was performed over a square ROI of average size 45 μm within the egg chamber. For each time delay Δ*t*, multiple of the delay Δ*t*_0_ = 10 *s* between consecutive frames, we calculated the image structure function 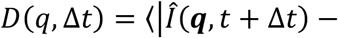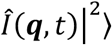, where 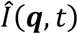 is the 2D Fourier transform of the image at time *t*, ***q*** is the wave vector, *q* = |***q***|, and the symbol 〈∙〉 indicates a combined temporal and azimuthal (*i.e.* performed over the orientation of ***q***) average. The obtained image structure function can be written as

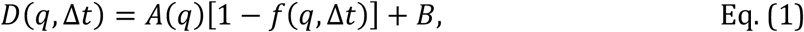

where *A*(*q*) is an amplitude term determined by the optical properties of the sample and by the microscope collection optics, *B* is a term accounting for the delta-correlated noise in the detection chain, and *f*(*q*, Δ*t*) is the intermediate scattering function (ISF)^79^, whose decay with Δ*t* mirrors the motion of the moving entities in the image.

As first step of our DDM analysis, we estimated the noise term *B* from an exponential fit of the tail of *D*(*q*, Δ*t*_0_) for *q* > 12 *μm*^−1^ (**Supplementary Fig. 5d**), a procedure that exploits the fact that *A*(*q*) vanishes for large enough *q* due to the finite numerical aperture of the microscope objective. Once *B* was known, we proceeded with the determination of the amplitude *A*(*q*) as the large Δ*t* limit of *D*(*q*, Δ*t*), a procedure that exploits the fact that *f*(*q*, Δ*t*) decays almost completely to zero for large Δ*t*. In our experiments, for which this typically happens for *q* > 0.4 *μm*^−1^, we estimated *A*(*q*) from a stretched exponential fit of *D*(*q*, Δ*t*) for Δ*t* > 100 *s* (**Supplementary Fig. 5d**).

Once *A* and *B* were known, Eq. (1) was inverted to obtain the ISF *f*(*q*, Δ*t*), as shown in **Supplementary Fig. 5e**. For identical particles with Gaussianly distributed displacements, the particle mean squared displacement can be obtained directly from *f*(*q*, Δ*t*) *via* a simple algebraical inversion. In less ideal cases, the following relation still holds34:

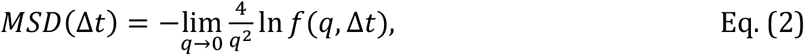

where *MSD*(Δ*t*) is a mean square displacement averaged over the entire population of particles present in the image. In our case, the limit for *q* → 0 was calculated via a linear extrapolation over the wave vector range 0.4 − 0.8 *μm*^−1^, as shown in the inset of **Supplementary Fig. 5e**. The results of this procedure, enabling the automatic determination of the average mean square displacement are shown in **Supplementary Fig. 5f** for all the egg chambers considered in this study. In all cases, the *MSD*(Δ*t*) displayed a linear dependence on Δ*t*, indicating a diffusive-like behavior with negligible persistence, at least at the investigated time scales. The corresponding effective diffusion coefficient *D* was estimated from the linear fit *MSD*(Δ*t*) = 4*D*Δ*t*. After averaging the obtained values over all examined cells of the same type, we found *D*_*WT*_ = 0.023 ± 0.006 *μm*^2^/*s* and *D*_*KO*_ = 0.021 ± 0.005 *μm*^2^/*s*, for wild-type and *Hecw^KO^*, respectively. These two values are fully compatible within the experimental uncertainty, indicating no significant difference in the RNP dynamics between wild-type and *Hecw^KO^* egg chambers.

### Statistical Analysis

Statistical analyses were performed with Prism (GraphPad software). Unless differently specified, all the statistical significance calculations were determined by using either unpaired Student’s t test or the non-parametric Mann Whitney test, after assessing the normal distribution of the sample with Normal (Gaussian) distribution test. Sample sizes were chosen arbitrarily with no inclusion and exclusion criteria. The investigators were not blind to the group allocation during the experiments and data analyses.

### Antibodies

Anti-Hecw was produced in rabbit by immunisation with the N-terminal fragment of Hecw (aa 1-130) protein fused to GST protein (Eurogentech S.A.). The resulting polyclonal antibody was affinity purified and validated (**Supplementary Fig. 1e**) on S2 cells depleted of Hecw according to the protocol described in^80^. For double-stranded RNA production, the following T7- and T3-tagged primers were used to amplify the *Hecw* region of interest:

Hecw^KD^ F: 5’-taatacgactcactatagggagaGGATAATTGCCACGATTGGT-3’,
Hecw^KD^ R: 5’-aattaaccctcactaaagggagaGGCGCCAATCGTTTGTG-3

All the other antibodies used in this study are listed in the following table.

| Antibody | Species | Supplier | code | WB | IF | IP |
| --- | --- | --- | --- | --- | --- | --- |
| anti-elav | rat | DSHB | 7E8A10 |  | 1:50 |  |
| anti-Hecw | rabbit | in house |  | 1:250 | 1:X00 | 4 µg/mg |
| anti-GST | rabbit | in house |  |  |  | 4 µg/mg |
| GFP-TRAP_A | Llama | Chromotek | gta-20 |  |  | 50 µl/mg |
| anti-GFP | rabbit | Sigma | G1544 | 1:5000 |  |  |
| anti-Fmrp | mouse | DSHB | 5A11-s | 1:300 | 1:300 | 3 µg/mg |
| anti-Grk | mouse | DSHB | 1D12-s | 1:400 | 1:200 |  |
| anti-K63 ubiquitin | rabbit | Millipore | clone Apu-3 | 1:500 | 1:250 |  |
| anti-K48 ubiquitin | rabbit | Millipore | clone Apu-2 | 1:500 | 1:250 |  |
| anti-Orb | mouse | DSHB | 4H8-s | 1:250 | 1:250 |  |
| anti-Orb | mouse | DSHB | 6H4-s | 1:250 | 1:250 | 1 µg/mg |
| Anti-Oskar | rabbit | Ephrussi lab | SA7299 |  | 1:2000 |  |
| anti-Ub | mouse | Enzo Life Science | clone FK2 |  | 1:300 | 20 µg/mg |
| anti-Ub | mouse | in house | ZTA10 | 1:5 |  |  |
| anti-alpha tubulin | rat | AbD Serotec | MCA78G |  | 1:100 |  |
| anti-tubulin | mouse | Sigma |  | 1:1000 |  |  |
| phalloidin-TRIC |  | Sigma | P1958 |  | 1:50 |  |
| anti-mouse IgG HRP | goat | Bio-Rad | 1721011 | 1:10000 |  |  |
| anti-rabbit IgG HRP | goat | Bio-Rad | 1706515 | 1:10000 |  |  |
| anti-rabbit Alexa647 | donkey | Thermo Fischer | A31571 |  | 1:400 |  |
| anti-rabbit Alexa488 | donkey | Thermo Fischer | A21206 |  | 1:400 |  |
| anti-rat Alexa647 | goat | Thermo Fischer | A21247 |  | 1:400 |  |
| anti-mouse Alexa488 | donkey | Thermo Fischer | A21202 |  | 1:400 |  |
| anti-rabbit Cy3 | donkey | Jackson Lab | 711-165-152 |  | 1:400 |  |
| anti-mouse Cy3 | donkey | Jackson Lab | 715-165-150 |  | 1:400 |  |

### Protein expression and purification

GST fusion proteins were expressed in Rosetta cells (Novagen) at 18°C for 16 hours after induction with 500 μM IPTG at an OD_600_ of 0.6. Cell pellets were resuspended in lysis buffer (50 mM Na-HEPES, pH 7.5, 200 mM NaCl, 1 mM EDTA, 0.1% NP40, 5% glycerol, and Protease Inhibitor Cocktail set III (Calbiochem)). Sonicated lysates were cleared by centrifugation at 20,000 rpm for 45 minutes. Supernatants were incubated with 1 ml of Glutathione SepharoseTM 4B beads (GE Healthcare) per liter of bacterial culture. After 4 hours at 4°C, beads were washed with PBS and equilibrated in maintenance buffer (50 mM Tris-HCl, pH 7.4, 100 mM NaCl, 1 mM EDTA, 1 mM DTT, 10% glycerol). The His-tagged E1 enzyme Uba1 and His-tagged E2 enzyme Ube2D3 (UBCH5c) used for *in vitro* ubiquitination were produced as described in81. His-tagged Fmrp was expressed in Rosetta at 18 °C for 16 hr after induction with 1 mM IPTG at an OD_600_ of 0.6. Cell pellets were resuspended in Buffer A (50 mM NaH_2_PO_4_ pH 7.8, 300 mM NaCl, 10 % glycerol, 10 mM imidazole and protease inhibitors) and lysed by sonication. Cell debris were removed by centrifugation and supernatants were incubated with 1 ml of HisPur Ni-NTA resin (Life Technologies), previously washed 3 times with Buffer A. After 3 hours at 4°C, beads were washed three times with Buffer A, Buffer A with 1 M NaCl, and Buffer A containing 20 mM imidazole, respectively. Untagged wild-type Ub (Sigma) was resuspended in maintenance buffer (50 mM Tris-HCl, pH 7.4, 100 mM NaCl, 1 mM EDTA, 1 mM DTT, 10% glycerol) and was purified using a Superdex 75 size exclusion chromatography column (GE Healthcare). Ub mutants K63R and K48R are from Boston Biochem.

### *In vitro* Ubiquitination

Reaction mixtures contained purified enzymes (20 nM E1, 250 nM purified Ube2D3, 250 nM GST-Hecw), 300 nM substrate (NiNTA bound His-Fmrp) and 1.25 μM Ub either wild-type or mutants in ubiquitination buffer (25 mM Tris-HCl, pH 7.6, 5 mM MgCl_2_, 100 mM NaCl, 0.2 μM DTT, 2 mM ATP) were incubated at 30°C for 60 minutes. Samples were then centrifuged in order to separate the pellet (containing the ubiquitinated substrate) from the supernatant containing the enzymes and the free Ub chains eventually produced. Pellets were washed four times in WASH buffer (50 mM Tris-HCl pH 7.4, 300 mM NaCl, 0,1 % Triton-X100, 5% glycerol, 1M UREA) before loading on SDS-PAGE gel. Detections were performed by immunoblotting using the anti-Ub antibody according to the protocol described in^81^. Membranes were stained with Coomassie after immunoblotting to show equal loading.

### Western blots and immunoprecipitations

*Drosophila* tissues, collected as described in^72^, were homogenized with a pestle and incubated for 20 minutes on ice in RIPA buffer (50 mM Tris-HCl, 150 mM NaCl, 1 mM EDTA, 1% Triton X-100, 1% sodium deoxycholate, and 0.1% SDS) supplemented with a protease inhibitor cocktail (CALBIOCHEM), clarified by centrifugation and analysed by immunoblotting.

For co-immunoprecipitation experiments (**Fig. 6a, Supplementary Fig. 7a**), *Drosophila* ovaries were lysed in JS buffer (Tris–HCl pH 7.6, 150 mM NaCl, 10% glycerol, 1.5 mM MgCl_2_, 0.1 M sodium pyrophosphate pH 7.5, 0.1 M PMSF, 0.5 M sodium vanadate pH 7.5 in Hepes, 0.5 M NaF) supplemented with protease inhibitors (Calbiochem). After extensive washes with JS buffer, beads were re-suspended in Laemmli-buffer and proteins analysed by SDS-PAGE and immunoblotting.

In case of drug treatments, fly ovaries were dissected in *Drosophila* S2 medium with 10 % of FBS (Euroclone) and incubated with 50 μM MG132 on an orbital shaker for 2 hours (**Supplementary Fig. 7e**) prior lysis.

For immunoprecipitation of ubiquitinated proteins, 40 μl of Agarose-TUBEs 2 (UM402, LifeSensors) previously equilibrated in JS buffer, were incubated overnight with 1 mg of ovary lysate on an orbital shaker at 4°C. After overnight incubation, the lysate was spin down and the supernatant was re-incubated with additional 30 μl of TUBEs for 1.5 hours at 4°C. Beads were then washed, re-suspended in Laemmli-buffer, loaded on SDS-PAGE and analysed by immunoblotting.

### LC-MS/MS analysis

For identification of Hecw interactors, a GST fusion construct encompassing the two WW domains of Hecw (636-834aa) was used. Briefly, 2μM of GST proteins were incubated with 1 mg of S2 lysate for 2 hours at 4°C in YY buffer (50 mM Na-HEPES pH 7.5, 150 mM NaCl, 1mM EDTA, 1mM EGTA, 10% glycerol, 1% triton-100). After four washes of the GST proteins with YY buffer, specifically bound proteins were resolved on 4-12% gel (Invitrogen) and detected by Coomassie staining. Samples were processed for MS analysis according to the STAGE-diging protocol described in^82^. Peptide mixtures were acidified with 100 μL of 0.1% formic acid (FA, Fluka) and eluted by the addition of 100 μL of 80% ACN, 0.1% FA, followed by a second elution with 100% CAN. All of the eluates peptides were dried in a Speed-Vac and resuspended in 15 μL of solvent A (2% ACN, 0.1% formic acid) and 4 μL were injected for each analysis on the Q-Exactive –HF mass spectrometer. Peptides separation was achieved with a linear gradient from 95% solvent A (2 % ACN, 0.1% formic acid) to 50% solvent B (80% acetonitrile, 0.1% formic acid) over 33 minutes and from 50% to 100% solvent B in 2 minutes at a constant flow rate of 0.25 μL/min, with a single run time of 45 minutes. MS data were acquired using a data-dependent top 12 method, the survey full scan MS spectra (300–1650 Th) were acquired in the Orbitrap with 60000 resolution, AGC target 3e6, IT 20 ms. For HCD spectra resolution was set to 15000, AGC target 1e5, IT 80 ms; normalised collision energy 28 and isolation width of 1.2 m/z. Raw data were processed using Proteome Discoverer (version 1.4.0.288, Thermo Fischer Scientific). MS^2^ spectra were searched with Mascot engine against an in-house *Drosophila Melanogaster* Db revised version (according to^83^). Scaffold (version Scaffold_4.3.4, Proteome Software Inc., Portland, OR) was used to validate MS/MS based peptide and protein identifications. Peptide identifications were accepted if they could be established at a probability greater than 95.0% by the Peptide Prophet algorithm^84^ with Scaffold delta-mass correction. Protein identifications were accepted if they could be established at a probability greater than 99.0% and contained at least 2 unique high confident peptides.

## Acknowledgements

We thank Sebastiano Pasqualato for DNA constructs, Ilaria Busi for initial generation of *Hecw^CI^* mutants and Stefano Confalonieri for reviewing the Mascot *Drosophila melanogaster* proteins database. We thank Paolo Soffientini, Federica Pisati, Valentina Dall’Olio and Laura Tizzoni at Cogentech facilities (Milan, Italy) for support in mass spectrometry, immunohistochemistry and qPCR analysis. We thank BDSC, DGRC, and DSHB for providing valuable *Drosophila* reagents to us and the community. This work was supported by the Italian Ministry of Education, Universities and Research (PRIN 20108MXN2J to S.P.); the CARIPLO foundation (2017-0746 to E.M.); the Associazione Italiana per Ricerca sul Cancro, (Investigator grant 2017-20661 to T. V., MFAG 2018-22083 to F.G.). Valentina Fajner’s work is supported by the Associazione Italiana per la Ricerca sul Cancro. V.F. was and S.S. is a PhD student of the European School of Molecular Medicine (SEMM).

## Author Contributions

Conceptualisation: V.F., El.M., T.V. and S.P; V.F. performed all the *Drosophila* experiments; El.M. generated biochemical data and supervised work by S.S. and V.F; A.O., Em.M., and D.P. aided in all the imaging acquisition and FLIP experiments; F.G. and R.C. analysed and quantitatively elaborated the live images; F.N. performed tunnel experiment; El.M. and T.V. contributed to planning and interpretation of data; S.P. coordinated the team, designed and supervised the project and wrote the paper with contributions from all authors.

## Supplementary Figure Legends

**Supplementary Figure 1.**
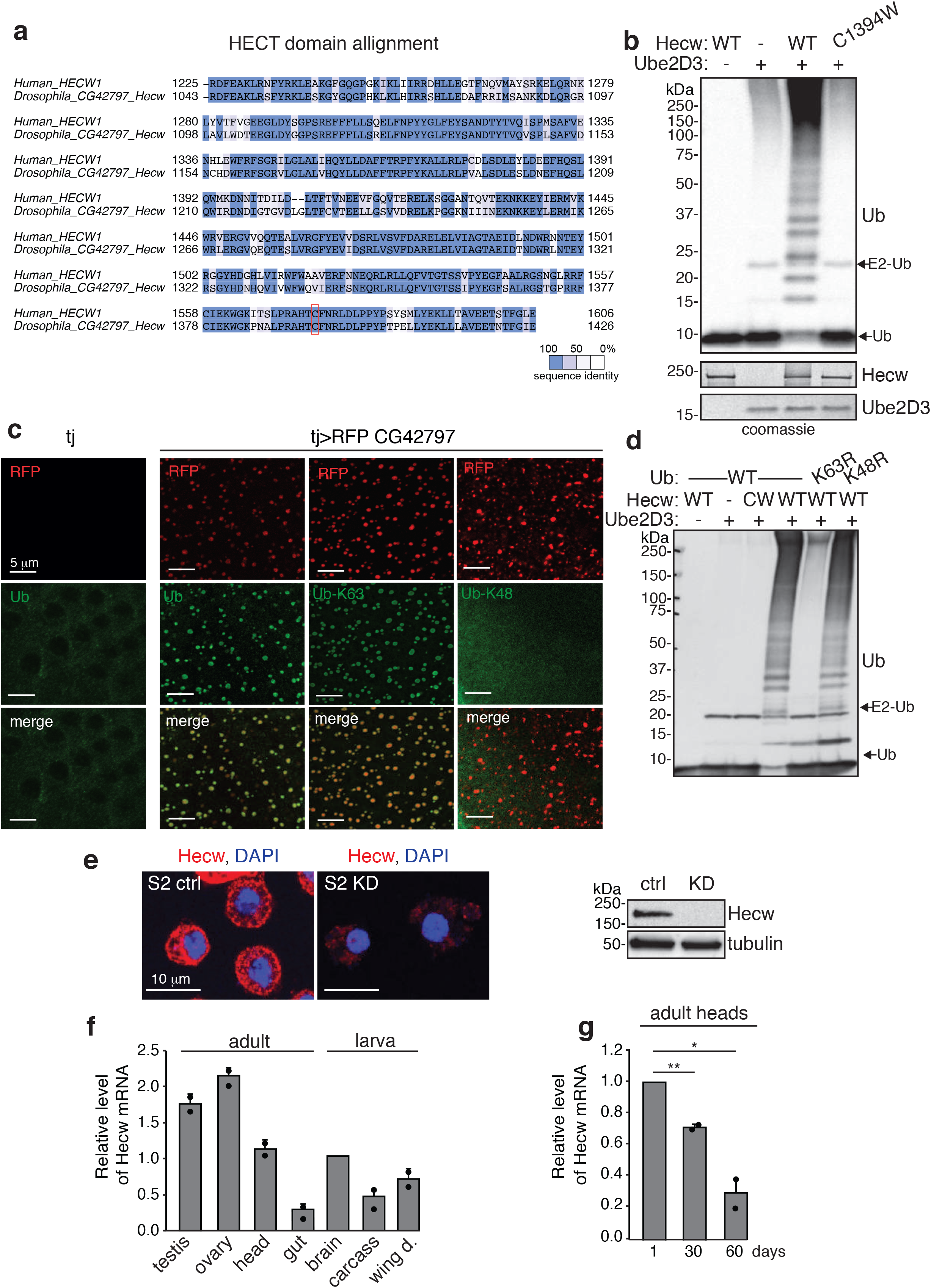
Hecw is a K63-specific ubiquitin E3 ligase. (**a**) Amino acid sequence alignment of the catalytic HECT domain of human HECW1 and *Drosophila* CG42797, color-coded according to sequence conservation. The catalytic cysteine is highlighted in the red box. ClustalW was used to create the sequence alignment of HECT between human and *Drosophila* proteins. (**b**) *In vitro* self-ubiquitination assay with the indicated recombinant proteins, analysed by IB with anti-Ub antibody. The reaction was quenched after 60 minutes. C1394W, a catalytic inactive mutant harbouring the same mutation present in *Hecw^CI^*. Coomassie shows equal loading (lower panel). (**c**) IF analysis of *Drosophila* follicle cells with the indicated fluorescent tag and antibodies. RFP-CG42797 is overexpressed in follicle cells with the *traffic jam-GAL4* driver (tj). RFP-Hecw localises in puncta that co-localises with a pan ubiquitin antibody (FK2, Ub) and an antibody that specifically recognises K63-linked chains, but not with an anti-K48 antibody. Left panels, control (tj driver only) shows no RFP signal and no accumulation of ubiquitinated puncta. Confocal images are shown. Scale bar: 5 μm. (**d**) *In vitro* self-ubiquitination assay with the indicated recombinant proteins and ubiquitin wild-type, K63R or K48R mutants. (**e**) IF analysis of *Drosophila* S2 cells untreated (ctrl) or depleted of Hecw by treatment with dsRNA (KD) for 48 hours. Hecw, red; DAPI, blue. Scale bar, 10 μm. Right panel, lysates from the same cells were IB as indicated. (**f**) *Hecw* mRNA expression measured in the indicated *Drosophila* tissues by qPCR. Expression levels are relative to larval brain and SD is calculated over two experiments with three technical replicates. (**g**) *Hecw* expression measured by qPCR in adult heads of the indicated age. Expression levels are relative to 1-day-old flies and SD is calculated over two experiments with three technical replicates.

**Supplementary Figure 2.**
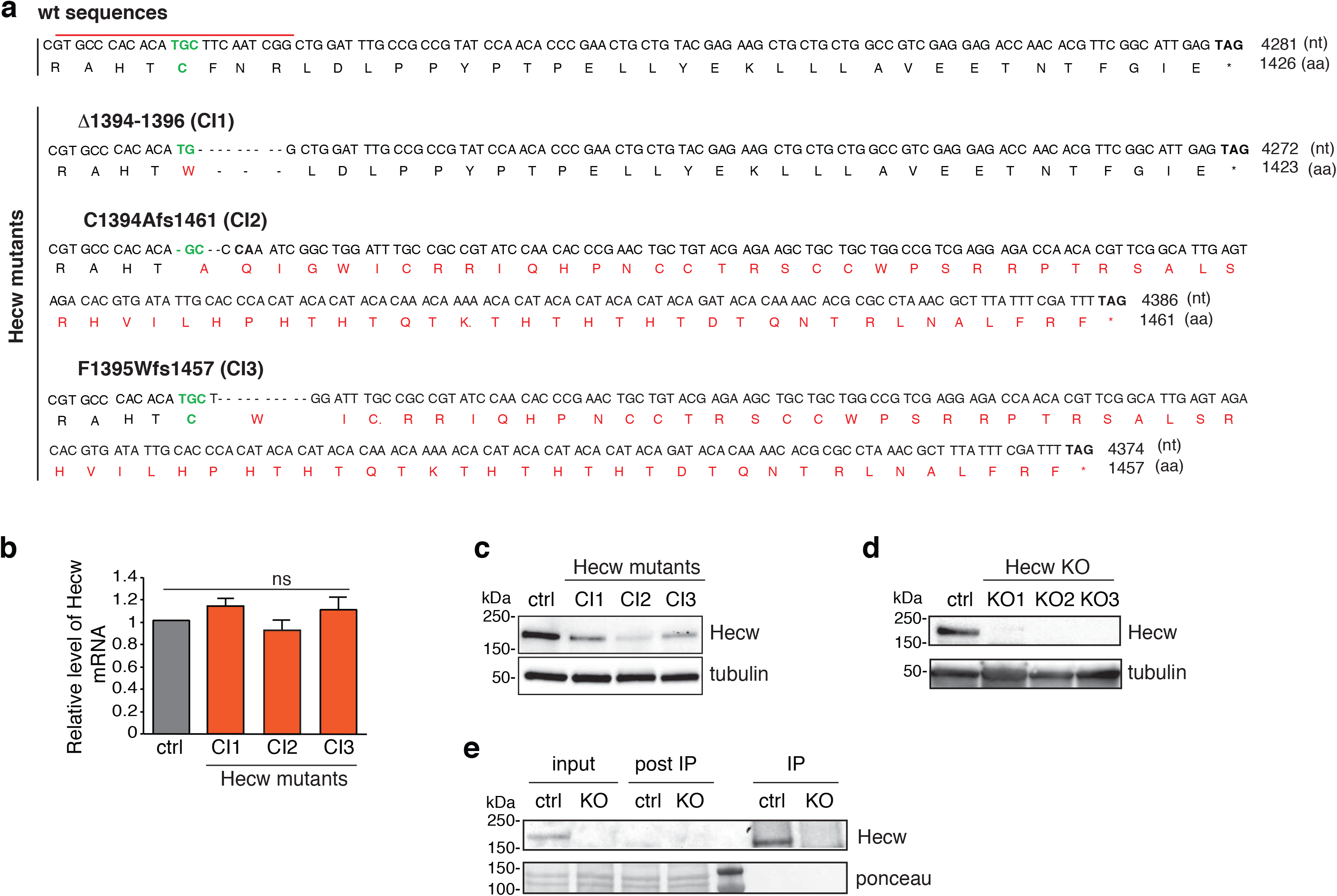
Characterisation of Hecw catalytic mutants generated by CRISPR/Cas9 technique. (**a**) Nucleotide and amino acid sequences of the mutants generated. Top panel: wild-type (wt) sequence of nucleotides (nt) and amino acids (aa). The catalytic cysteine in green and the stop codon is indicated with an asterisk. The red line indicates the region targeted by the Cas9 guide. In mutant sequences, deletions are indicated with dashes and amino acids that differ from the wild-type are in red. (**b**) mRNA levels of *Hecw* were measured by qPCR in larval brains of the mutated lines. No significant differences were scored. Expression levels are relative to wild-type larval brain and SD is calculated over three experiments with three technical replicates (**c**) IB analysis of larval brains of the mutated lines. The expression levels of Hecw mutant proteins are reduced in comparison with wild-type control (yw). (**d**) IB analysis of adult ovaries of the Hecw^KO^ lines. (**e**) 0.5 mg of ovary lysates from the indicated lines were IP with the anti-Hecw antibody and IB as indicated. Post IP, supernatant post IP.

**Supplementary Figure 3.**
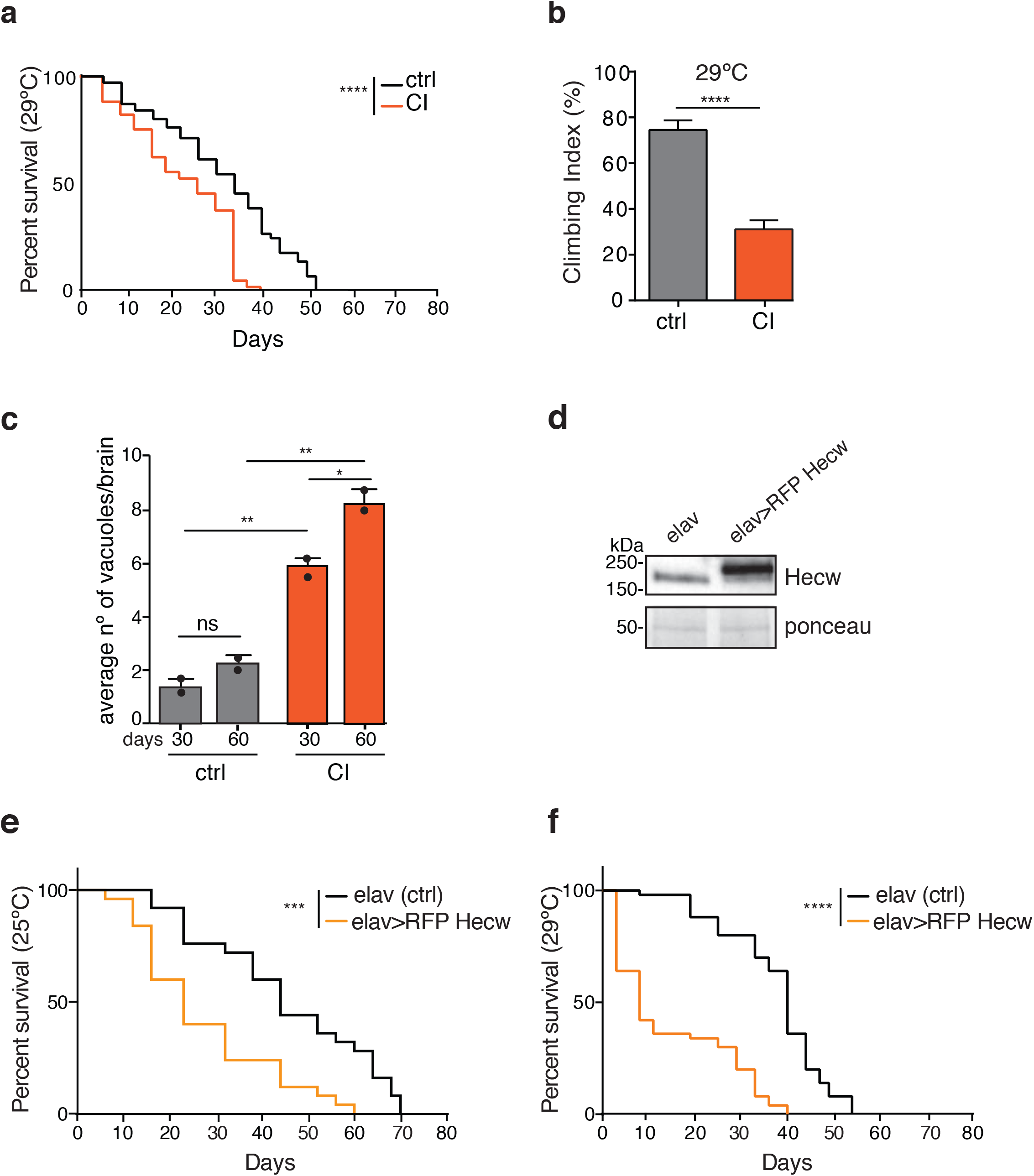
Hecw depletion and overexpression cause reduced longevity and premature motor function impairment. (**a**) Survival curve of the indicated genotypes. Percentage of survival was calculated over 100 animals at 29°C. *Hecw^CI^* (CI) shows a significant decrease in lifespan compared with the control line (****P<0,0001 with log-rank test). (**b**) Negative geotaxis assay performed on 12 day-old flies at 29°C. Results are expressed as a mean of 8 groups of animals (*n*=8/group) ± s.e.m. Mutant flies show a climbing deficit, ****P<0,0001 with Mann Whitney test. (**c**) Quantification of vacuoles with diameter >2 μm in 30-day-old fly brains of the indicated genotypes, *n*=5 animals/genotype. Results are expressed as mean of two biological replicates ± s.e.m. **P<0,01 with t test. (**d**) IB analysis of adult head lysates with the indicated antibodies. RFP-Hecw induced by the *elav-GAL4* driver (elav> RFP Hecw) runs at higher mw in comparison with wild-type Hecw protein expressed in the control line (elav driver only). (**e,f**) Survival curve of the indicated genotype at 25°C (**e**) and 29°C (**f**). Percentage of survival was calculated over 50 animals per genotypes. Flies with overexpressed Hecw in neurons show a significant decrease in their lifespan compared with the control line (**** P<0,001 with log-rank test).

**Supplementary Figure 4.**
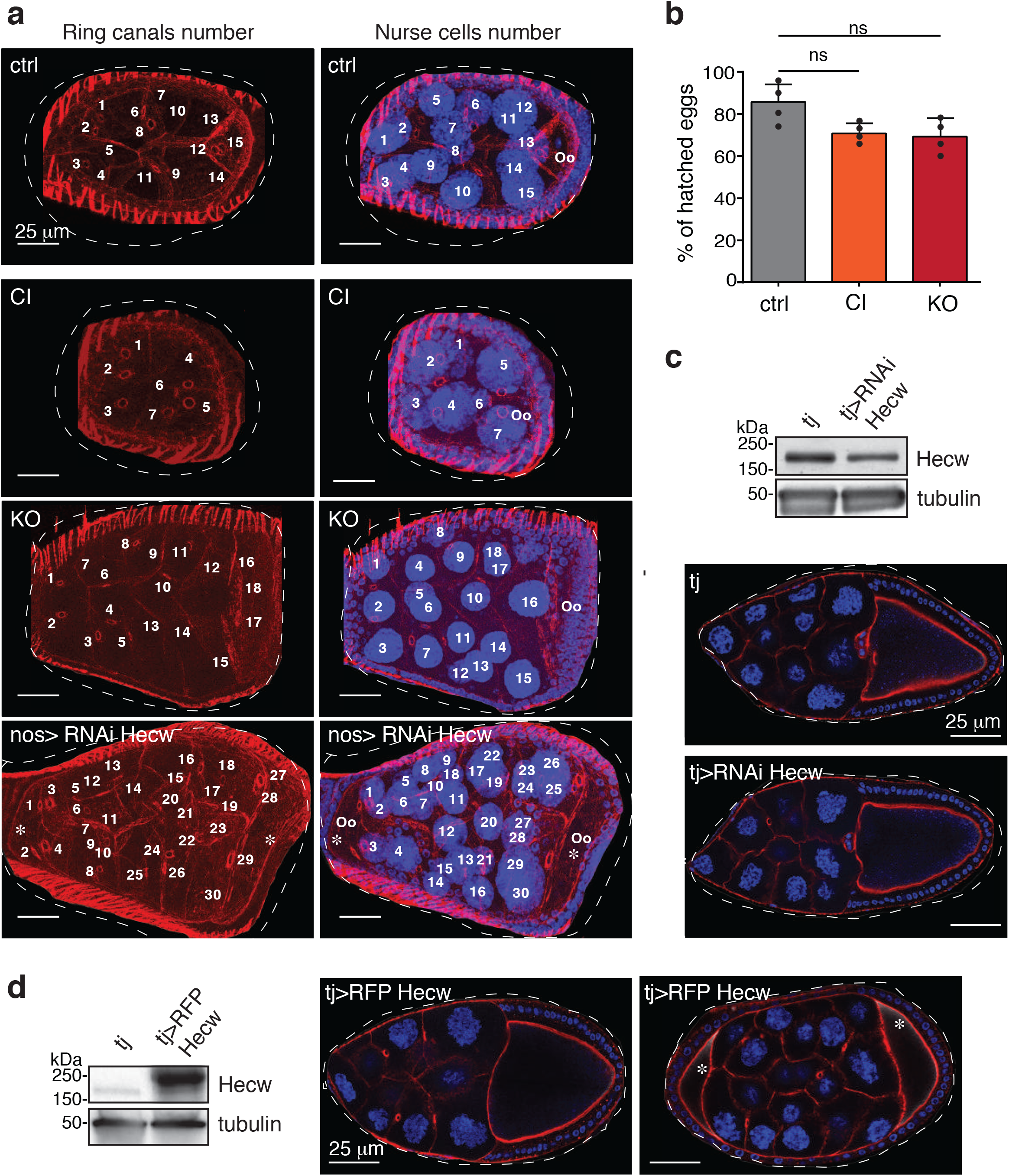
Defective egg chambers upon Hecw perturbation. (**a**) Confocal analysis of ovaries of the indicated genotypes. Maximal projection of the z-stack images is shown. Total number of nurse cells and ring canals were counted to assess the amount of germline cells and cell division. Observed defects include a reduced number of nurse cells (CI, *Hecw^CI^* mutant is shown as an example), an increased number of nurse cells (KO, *Hecw^KO^* is shown as an example), and compound egg chambers (Hecw germline-specific knock down *nos>RNAi Hecw* is shown as an example; white asterisks indicate the two oocytes). Phallodin, red; DAPI, blue. Scale bar: 25 μm. (**b**) Hatching rate of eggs laid in 24 hours by 20-day-old flies of the indicated genotypes. Results are expressed as mean of number of laid eggs ± SD calculated over four experiments (*n*=6 females/exp). (**c**) IB analysis of ovary lysates from control (follicle cells driver only, *traffic jam-GAL4* tj) and Hecw knock down (*tj>UAS-RNAi Hecw*) 3-day-old flies. Note that depletion appears limited as follicle cells represents 20-25% of the single egg chamber. Bottom, IF analysis of the same egg chambers. Red, phalloidin; blue, DAPI. Scale bar: 25 μm. Ovaries with Hecw depletion only in follicle cells do not present oogenesis defects. (**d**) IB and IF analysis of the indicated genotype, as in c. Most of the egg chambers with ectopic Hecw expression in follicle cells (tj>UAS RFP Hecw) do not present oogenesis defects (central panel) except for 3,5% of them that present compound egg chambers (right panel; white asterisks indicate the two oocytes). See also Supplementary Table 1 for the quantification.

**Supplementary Figure 5.**
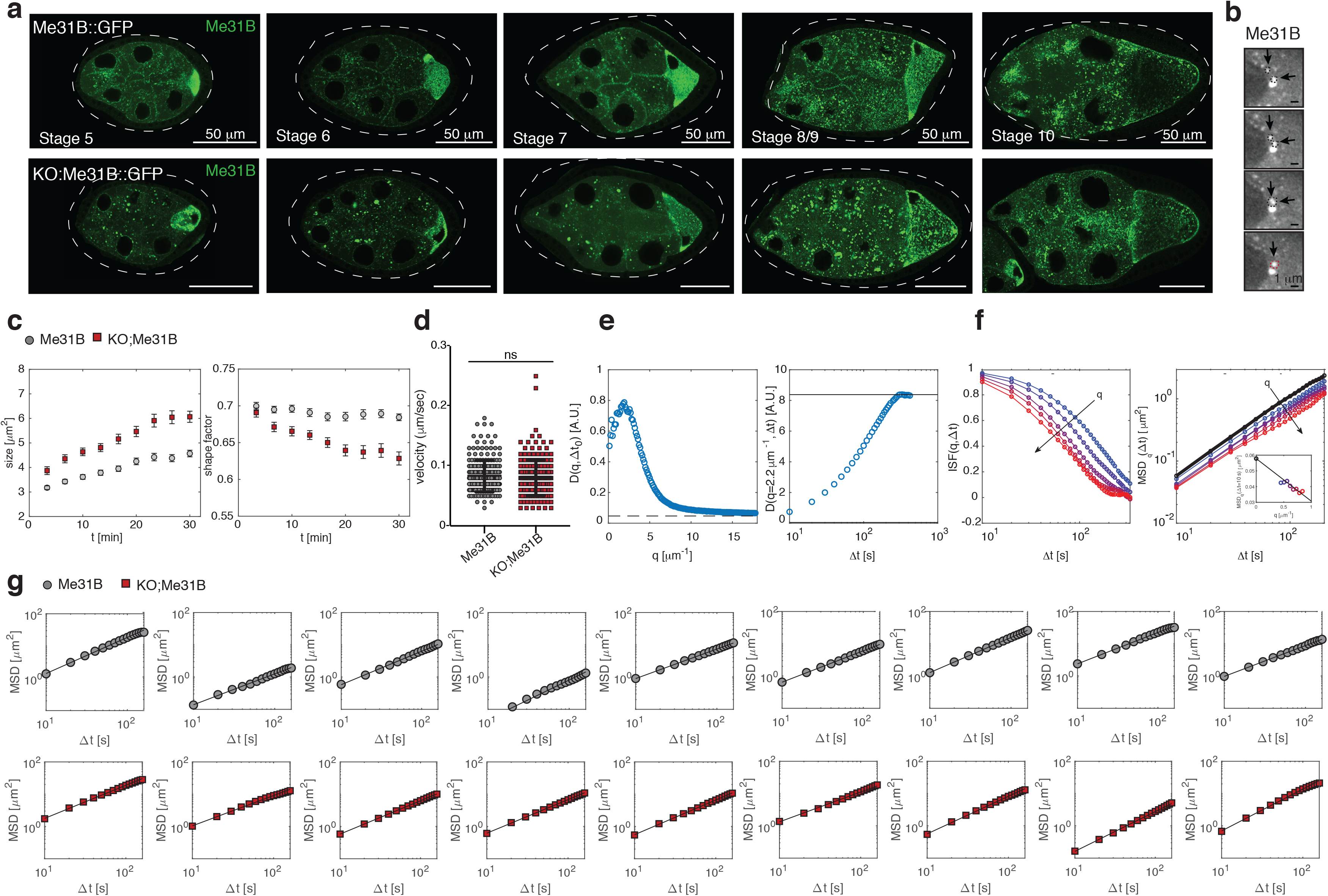
Mobility and coarsening of RNPs. (**a**) Examples of egg chambers from 3-day-old flies at different stages of development. Egg chambers of *Hecw^KO^; Me31B::GFP* animals show enlarged RNPs. Scale bar: 50 μm. (**b**) Fusion of RNPs (indicated by arrow heads and dashed circles in sequential frames) in *Me31B::GFP* egg chambers. Scale bar 1 μm. (**c**) Temporal evolution of size (left panel) and shape factor 4*πA*/*P*^2^ (right panel) of the RNPs during *ex-vivo* observation. A progressive increase in the average size of the RNPs (coarsening) is observed both in wild-type (grey dot, *Me31B::GFP*) and *Hecw^KO^* (red squares, *Hecw^KO^; Me31B::GFP*) flies ovaries. In wild-type egg chambers, the coarsening process takes place without any significant change in the geometry of the granules, which grow while maintaining their approximately spherical shape. On the contrary, in *Hecw^KO^* egg chambers, the increase of the average size of the granules is accompanied by a marked decrease of the shape factor, indicating the formation of irregularly shaped aggregates. (**d**) Quantification of the speed of tracked single RNP by live imaging analysis. The velocity of control (Me31B::GFP) and *Hecw^KO^* (KO;Me31B) particles is not statistically different (*v_wt_* = 0,0829 ± 0,026 *μ*m/s, *v_ko_* = 0,0804 ± 0,0298 *μ*m/s. P=0,3302, ns by Mann Whitney test). Statistical analysis was performed over 3 biological replicates for a total of n=220 particles for the control and n=213 for *Hecw^KO^* (11 egg chambers/sample). (**e**) Left: representative image structure function *D*(*q*, *Δt*) obtained from DDM analysis as a function of the wavevector *q* for fixed Δ*t* = Δ*t*_0_ = 10 *s*. For large *q* the function attains a plateau value (dashed horizontal line) corresponding to the noise contribution *B* in Eq. (1). For each value of *q*, the amplitude term *A*(*q*) is estimated from the large Δ*t* limit of *D*(*q*, *Δt*) (continuous horizontal line in the right panel). (**f**) Coloured empty symbols: intermediate scattering functions *f*(*q*, Δ*t*) for different *q* in the range 0.4 − 0.8 *μm*^−1^ (left) and corresponding *q*^2^dependent estimate of the mean square displacement 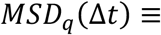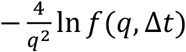 (right). Black symbols in the right panel corresponds to the best estimate for the average mean square displacement *MSD*(Δ*t*), obtained by extrapolating *MSD*_q_(Δ*t*) to *q* = 0 at fixed Δ*t*, as shown in the inset for Δ*t* = 10 *s*. (**g**) Average mean square displacement *MSD*(Δ*t*) of RNPs determined with Differential Dynamic Microscopy (DDM). Each panel corresponds to a different cell. The effective diffusion coefficients obtained by fitting a linear model to the data (*D*_*WT*_ = 0.023 ± 0.006 *μm*^2^/*s* and *D*_*KO*_ = 0.021 ± 0.005 *μm*^2^/*s*) show no significant difference in the RNP dynamics between for wild type (grey dot, Me31B) and *Hecw^KO^* egg chambers(red squares, KO;Me31B).

**Supplementary Figure 6.**
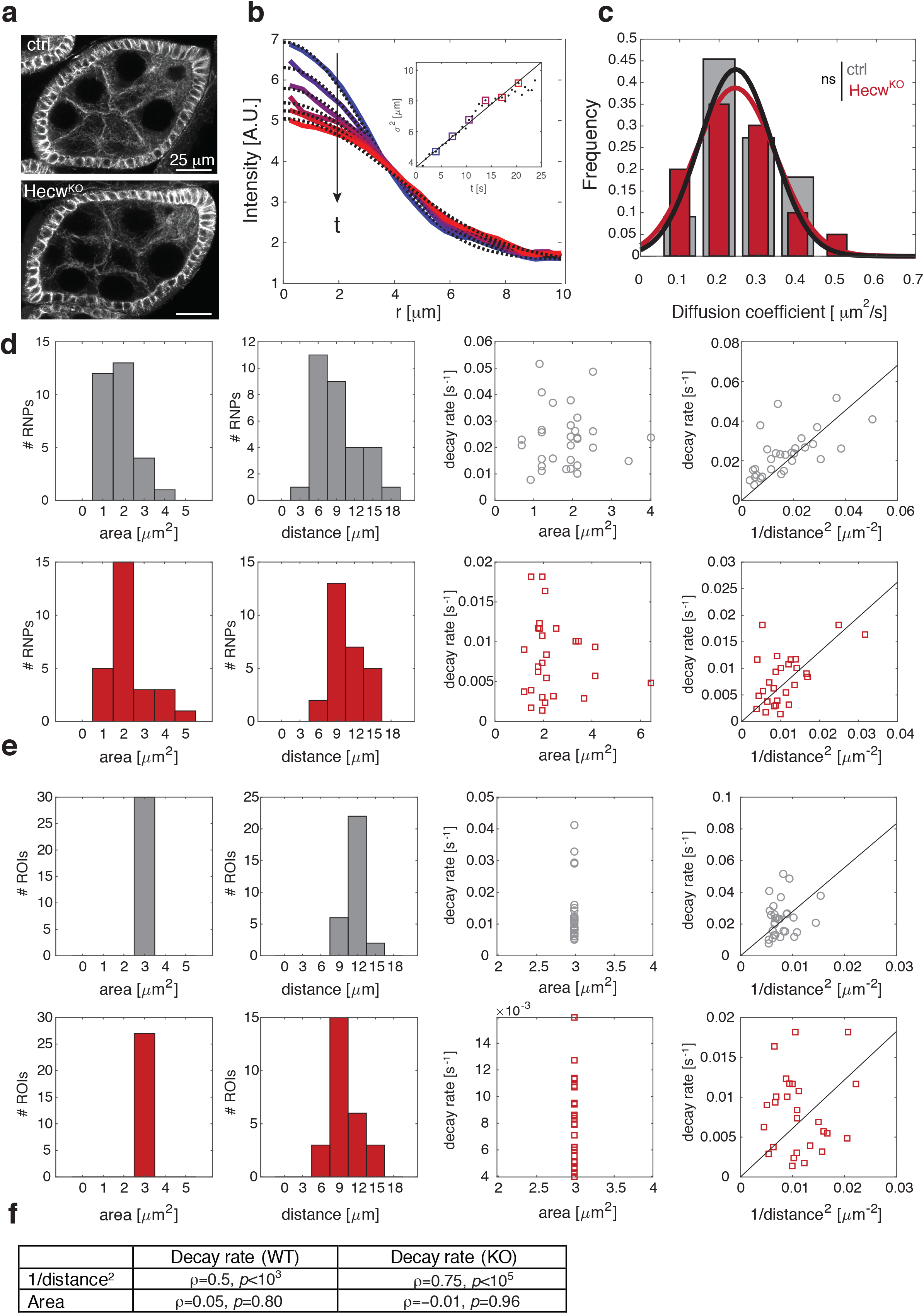
Free diffusion and microtubule-dependent movement of Me31B are not affected by the absence of Hecw. (**a**) IF analysis to detect the microtubule marker α-tubulin in control (*Me31B::GFP*) and *Hecw^KO^* (*Hecw^KO^1;Me31B::GFP*) egg chambers dissected from 3 day-old flies. (**b**) Continuous lines: azimuthally averaged intensity profiles measured at different times after photobleaching; dotted lines: best fitting Gaussian curves, enabling the estimation of a time-dependent width *σ*(*t*). In the inset, *σ*^2^(*t*) is plotted *vs* time. A linear fit to the data (continuous line) enables estimating the diffusion coefficient *D*. (**c**) Bars: frequency distributions of the obtained diffusion coefficients, showing no statistically significant differences between *Hecw^KO^* and wild-type egg chambers. Continuous lines are Gaussian fits to the data. (**d,e**) Analysis of the size and spatial distribution of RNPs (first and second rows) and cytoplasm ROIs (third and fourth rows) considered in the FLIP experiment. From left to right in each row: frequency distribution of the areas, frequency distribution of the distances from the centre of the photobleached area, estimated decay rate *vs* area, estimated decay rate *vs* inverse squared distance. Grey bars and symbols refer to control, red bars and symbols refer to *Hecw^KO^* cells. While in *Hecw^KO^* egg chambers, RNPs are, on average, larger than those in wild-type egg chambers, no statistically significant correlation is observed between size and decay time within each group of nurse cells. Within each group of cells, there is a significant (negative) correlation between the distance from the bleached area and decay time. The distributions of distances in the two groups are not significantly different (second column of panel d). (**f**) Table reporting the Pearson correlation coefficient between fluorescence decay rate of the RNPs and inverse of the squared distance from the center of the photobleached area (first row) and between decay rate and area (second row). The corresponding p-values are also reported.

**Supplementary Figure 7.**
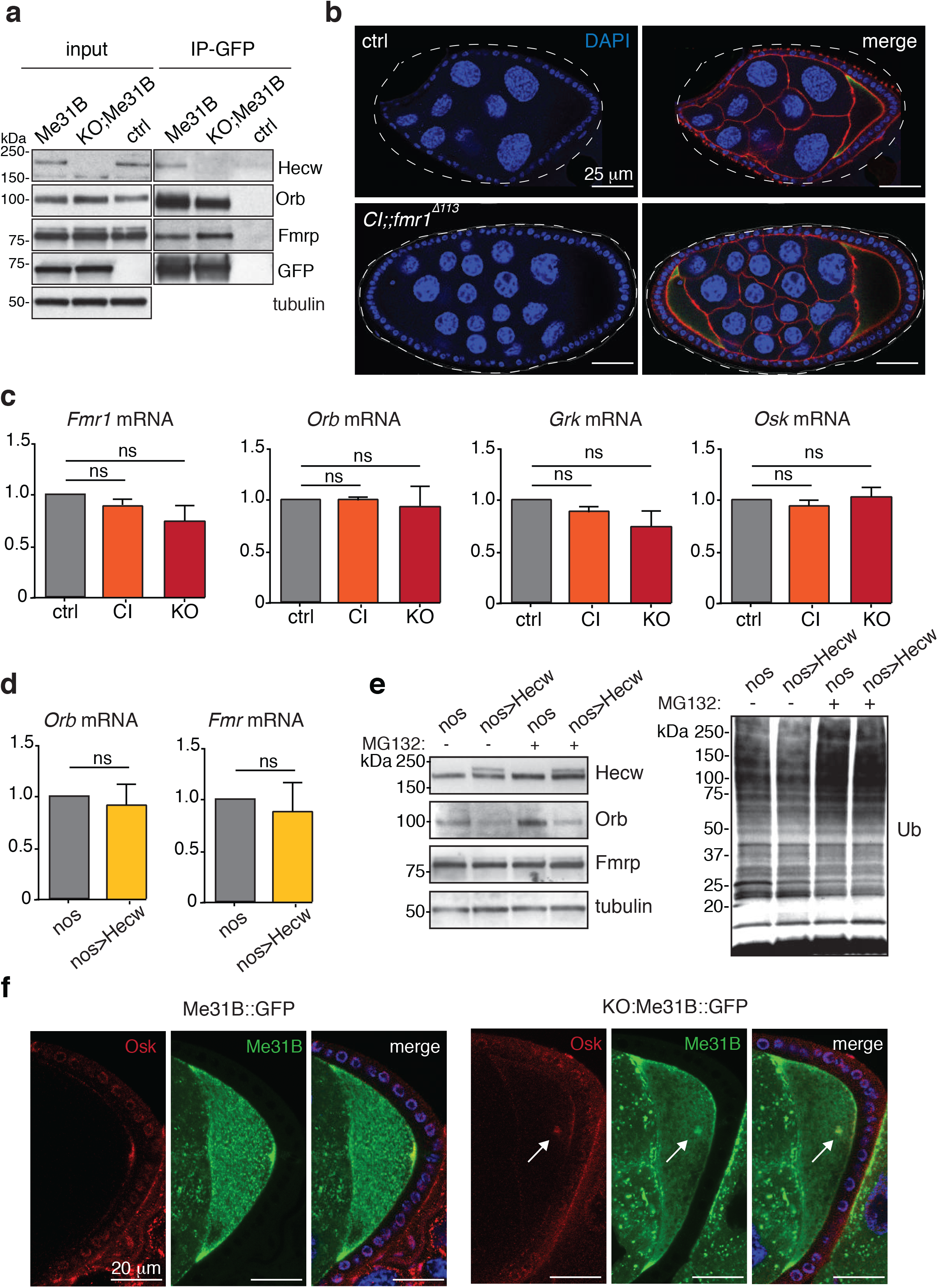
Hecw interacts with RNP components. (**a**) 0.5 mg of ovary lysates from the indicated lines were IP and IB as indicated. (**b**) IF analysis of egg chambers of 3-day-old control (yw) and *Hecw^CI^;;Fmr1 Δ113* double mutant (CI;;Fmr1) animals. Double mutant shows no worsening of the single mutant phenotypes. Compound egg chamber is reported in the figure as an example. (**c**) mRNA levels of *Fmr1, orb*, *grk* and *osk* was measured by qPCR in the adult ovaries of the indicated genotypes. The reported expression levels are relative to control and SD is calculated over two experiments with three technical replicates/each. (**d**) mRNA levels of *Fmr1* and *orb* in the adult ovaries of the indicated genotypes, measured by qPCR as in c. (**e**) IB of control (driver only, nos) and RFP-Hecw overexpressing fly ovaries (nos>RFP Hecw) treated with 50 μM MG132 for 2 hours, and probed with the indicated antibodies. Left panel: Orb level decreases upon Hecw overexpression in the germline and does not increase upon proteasome inhibition. Right panel: IB with anti-Ub antibody shows Ub accumulation upon MG132 treatment. (**f**) IF analysis of 3-day-old wild-type and *Hecw^KO^* egg chambers. Green, Me31B::GFP marks RNPs. Upper panels: red, Osk localises at the posterior margin of the oocyte in wild-type control. Bottom panels: white arrows indicate mislocalised Osk in Hecw mutant flies. Blue, DAPI.

**Supplementary Table 1.**
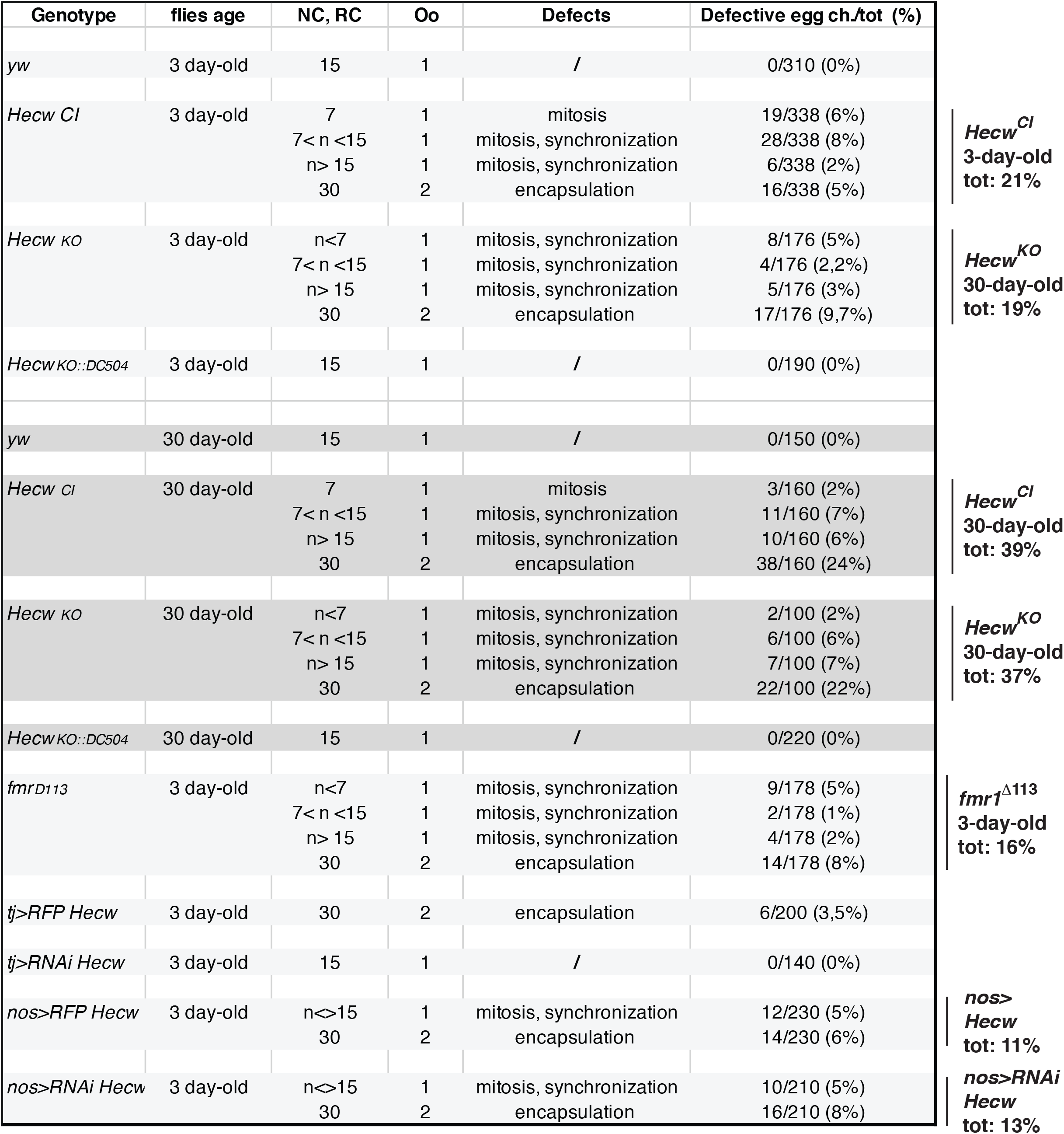
Classification of oogenesis defects. Complete classification of the defects observed in the indicated genotypes at the indicated time points. Numbers of nurse cells (NC), ring canals (RC), and oocytes (Oo) are specified in the third and fourth column, respectively.

**Supplementary Table 2.**
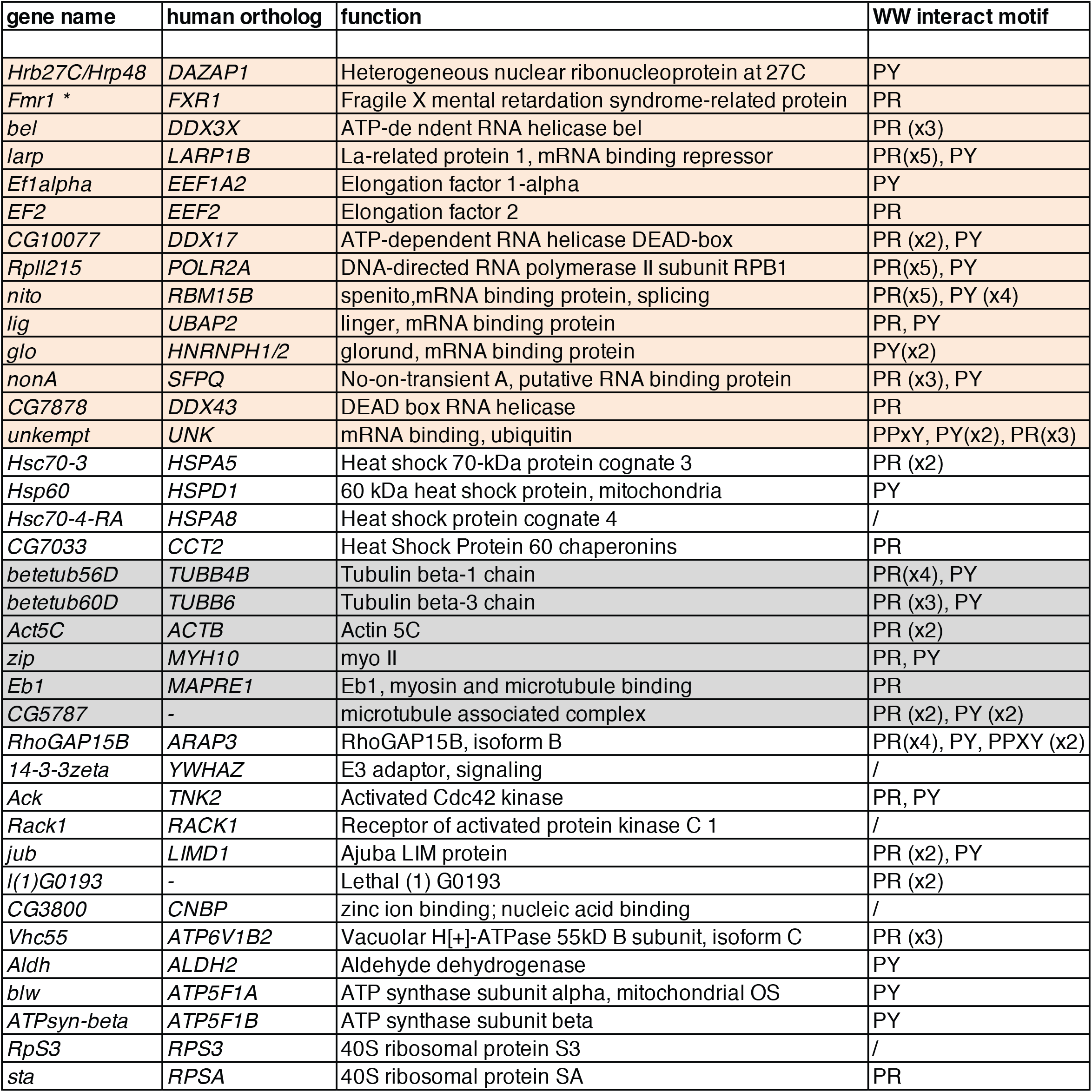
Hecw interactor candidates. List of Hecw interactors identified by mass spectrometry analysis.

**Supplementary Table 3.** Genotypes of strains used in figures.

| ***Drosophila* strains** | **Figure** | **Supplementary Figure** |
| --- | --- | --- |
| *Dp(1;3)DC504* | 3d |  |
| *elav-GAL4* |  | 3d-f |
| *elav-GAL4;;UAS-RFPHecw* |  | 3d-f |
| *fmr1*<sup>Δ113M</sup> | 6b |  |
| *Hecw*<sup>CI</sup> | 2a,c-e; 3a,b; 6b,f,h,i | 4a,b; 7c |
| *Hecw*<sup>CI</sup>;;*fmr* Δ113 |  | 7b |
| *Hecw*<sup>KO</sup> | 2b-f; 3a,b,d; 4a; 6e,f | 4a,b; 7c |
| *Hecw*<sup>KO;;DC504</sup> | 2d,e,g; 3d |  |
| *Hecw*<sup>KO</sup>;;*Me31B::GFP* | 4b-f; 5a-c; 6a | 5a-e; 6a,f; 7a,f |
| *Me31B::GFP* | 4b-f; 5a-c; 6a | 5a-e; 6a,f; 7a,f |
| *nanosGAL4-VP16* | 3c,e; 6g | 7d,e |
| *nanosGAL4-VP16;;UAS-HecwRNAi* | 3c | 4a |
| *nanosGAL4-VP16;;UAS-RFPHecw* | 3e; 6g | 7d,e |
| *traffic jam-GAL4* |  | 1,c; 4c,d |
| *traffic jam-GAL4; UAS-HecwRNAi* |  | 4c |
| *traffic jam-GAL4; UAS-RFPHecw* |  | 1,c; 4d |
| *y<sup>1</sup>w<sup>1</sup>* | 1b-d; 2a-g; 2a,b; 3a,b; 4a; 6a-c,e,f,h,i | 1f,g; 2b-e; 3a-c; 4a,b |

